# The Folding Pathway of an Ig Domain is Conserved On and Off the Ribosome

**DOI:** 10.1101/253013

**Authors:** Pengfei Tian, Annette Steward, Renuka Kudva, Ting Su, Patrick J. Shilling, Adrian A. Nickson, Jeffrey J. Hollins, Roland Beckmann, Gunnar von Heijne, Jane Clarke, Robert B. Best

## Abstract

Proteins that fold cotranslationally may do so in a restricted configurational space, due to the volume occupied by the ribosome. How does this environment, coupled with the close proximity of the ribosome, affect the folding pathway of a protein? Previous studies have shown that the cotranslational folding process for many proteins, including small, single domains, is directly affected by the ribosome. Here, we investigate the cotranslational folding of an all-b immunoglobulin domain, titin I27. Using an arrest peptide-based assay and structural studies by cryo-EM, we show that I27 folds in the mouth of the ribosome exit tunnel. Simulations that use a kinetic model for the force-dependence of escape from arrest, accurately predict the fraction of folded protein as a function of length. We used these simulations to probe the folding pathway on and off the ribosome. Our simulations - which also reproduce experiments on mutant forms of I27 - show that I27 folds, while still sequestered in the mouth of the ribosome exit tunnel, by essentially the same pathway as free I27, with only subtle shifts of critical contacts from the C to the N terminus.

**Significance Statement:** Most proteins need to fold into a specific three-dimensional structure in order to function. The mechanism by which isolated proteins fold has been thoroughly studied by experiment and theory. However, in the cell proteins do not fold in isolation, but are synthesized as linear chains by the ribosome during translation. It is therefore natural to ask at which point during synthesis proteins fold, and whether this differs from the folding of isolated protein molecules. By studying folding of a well characterized protein domain, titin I27, stalled at different points during translation, we show that it already folds in the mouth of the ribosome exit tunnel, and that the mechanism is almost identical to that of the isolated protein.

## Introduction

To what extent is the cotranslational folding pathway of a protein influenced by the presence of the ribosome and by the vectorial emergence of the polypeptide chain during translation? Recent studies have shown that small proteins can fold inside the ribosome exit tunnel (e.g., the small zinc finger domain ADR1a) (1), while other proteins can fold at the mouth of the tunnel (e.g., the three-helix bundle spectrin domains) (2); however some proteins may be simply too large to fold within the confines of the ribosome (e.g., DHFR) (3). The nature of cotranslational protein folding is determined by a number of biophysical factors, including the folding properties of the isolated protein (4–9), together with the effects the ribosome itself may have on the folding process (10–16). Due to the spatial constraints imposed upon the nascent chain by the confines of the tunnel, and effects due to the close proximity of the ribosome itself, the ribosome has been shown to influence directly the cotranslational folding of small proteins and single domains: The stability of folded or partly folded states may be reduced when folding occurs close to, or within the confines of, the ribosome (17, 18); the folding kinetics are expected to be correspondingly altered, with the rate of folding likely to be decreased and the unfolding rate increased, in close proximity to the ribosome(18). Interactions of the folded state or nascent polypeptide with the ribosome may also be either stabilising or destabilising (19, 20). Since translation is vectorial in nature, it is possible that when proteins fold cotranslationally they fold via different pathways than those used when proteins fold outside the ribosome, or when isolated proteins fold *in vitro* (2, 11, 21–24). However, addressing these issues is challenging, because standard protein folding methods are not directly applicable to cotranslational folding.

The folding of the protein close to the ribosome generates a pulling force on the nascent chain. This force has been probed by single molecule (25) as well as arrest peptide (AP) experiments (1–3). In this work, we use such arrest peptide-based cotranslational force-measurement experiments, simulations, and structural studies to investigate how the ribosome affects the folding of titin I27, a small all-β immunoglobulin domain with a complex greek-key fold; the stability, kinetics and folding pathway of I27 has been extensively characterized in previous studies of the isolated domain (26, 27). In this study we investigate whether I27 can begin to fold in the confines of the ribosome, and if the folding pathway observed in the isolated domain is conserved during cotranslational folding. Results from all three techniques show that I27 folds in the mouth of the ribosome exit tunnel; our simulations correctly capture the onset of folding in I27 and three mutant variants, allowing us to predict how destabilisation of regions that fold early and late in the isolated domain affect folding on the ribosome. Our simulations further show that the folding pathway of I27 is largely unaffected by the presence of the ribosome, except for small but significant changes observed for contacts near the N and C termini.

## Results

### I27 folds close to the ribosome

In order to gain insight into when I27 can commence folding on the ribosome, we employed an arrest peptide force-measurement assay (28) carried out using the PURE *in vitro* translation system, as described in (1–3). In these experiments, the *E. coli* SecM arrest peptide (AP) is used to stall the nascent protein chain temporarily during translation. The yield of full-length protein which escapes stalling in a defined time interval (*f*_*FL*_), determined from SDS-PAGE gels, provides a proxy for the pulling force exerted on the nascent chain by the protein as it folds (1–3) (Figure 1A). By measuring *f*_*FL*_ for a set of constructs where the length *L* of the linker between the target protein and the SecM AP is systematically varied, a force profile can be recorded that reflects the points during translation where the folding process starts and ends. Previous work has shown that the location of the main peak in a force profile correlates with the acquisition of protease resistance in an on-ribosome pulse-proteolysis assay (17, 29) and that the amplitude of the main force peak correlates with the thermodynamic stability of the protein (29, 30), indicating that the main peak represents a *bona fide* folding event rather than, *e.g.*, the formation of a molten globule. The sharp onset of the main force peak observed for most proteins analysed thus far (29) is also as expected for a cooperative folding event.

The force profile for wild-type I27 (Figure 1B) has a distinct peak at *L* = 35-38 residues (see Methods for sequences of the constructs). This peak is absent from the force profile for the mutant I27[W34E], a non-folding variant of I27, demonstrating that the peak is due to a folding event and not, for example, to non-specific interactions of the unfolded nascent chain with the ribosome. The non-zero *f*_*FL*_ for the non-folding mutant is attributed to the spontaneous rate of escape from arrest in the absence of acceleration by forces associated with folding. Since it takes ~35 residues in an extended conformation to span the ~100 Å long exit tunnel (31), the critical length *L* ≈ 35 residues suggests that I27 starts folding while in mouth of the exit tunnel.

**Figure 1.**
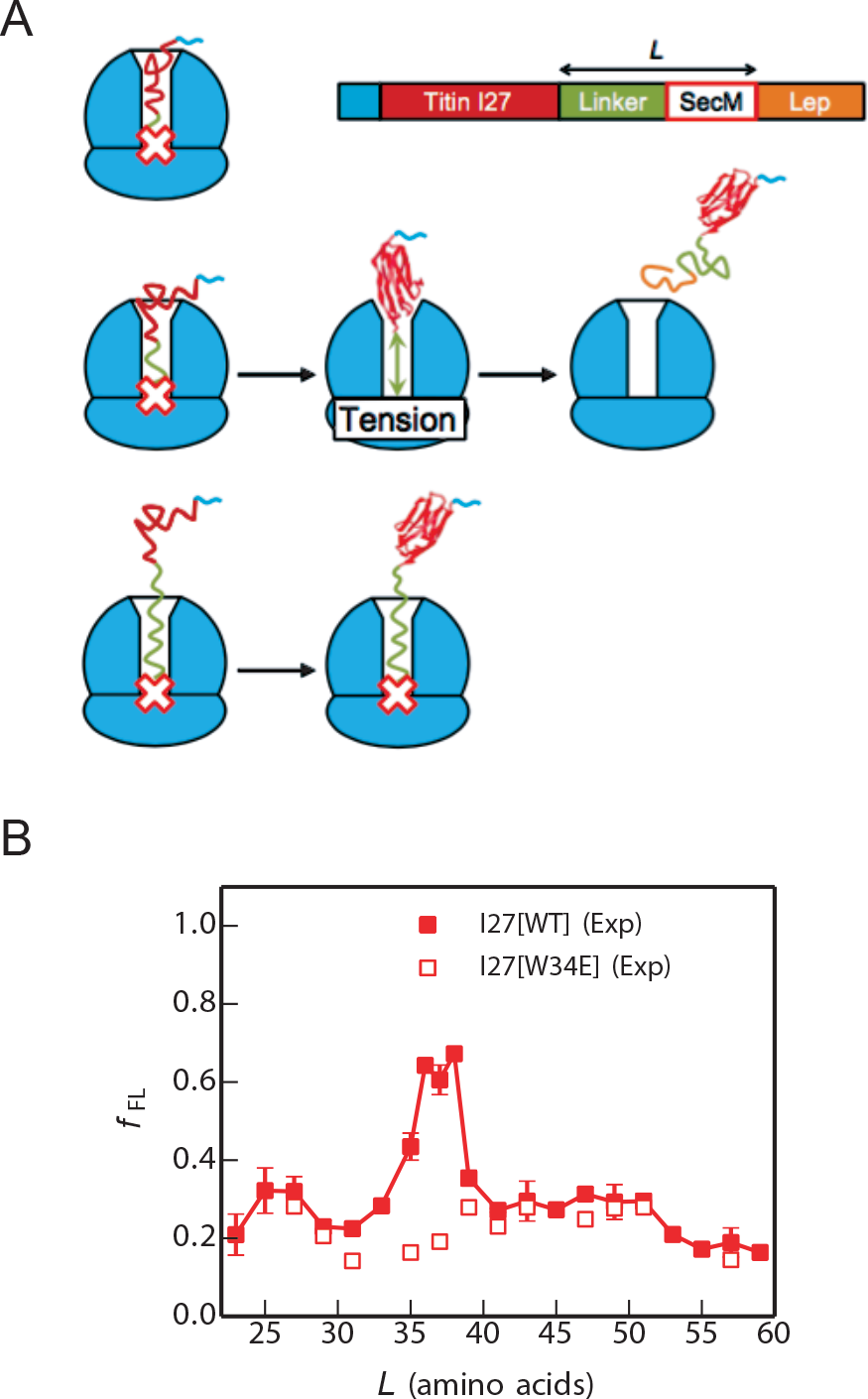
Cotranslational folding of the titin I27 domain by force-profile analysis. (A) The force-measurement assay (modified from (2)). I27, preceded by a His-tag, is placed *L* residues away from the last amino acid of the SecM AP, which in turn is followed by a 23-residue C-terminal tail derived from *E. coli* LepB. Constructs are translated for 15 min. in the PURE *in vitro* translation system, and the relative amounts of arrested and full-length peptide chains produced are determined by SDS-PAGE. The fraction full-length protein *f*_*FL*_, reflects the force exerted on the AP by the folding of I27 at linker length *L*. At short linker lengths (top), there is not enough room in the exit tunnel for I27 to fold, little force is exerted on the AP, and the ribosome stalls efficiently on the AP (*f*_*FL*_ ≈ 0). At intermediate linker lengths (middle), there is enough room for I27 to fold but only if the linker segment is stretched, force is exerted on the AP, and stalling is reduced (*f*_*FL*_ > 0). At long linker lengths (bottom), I27 has already folded when the ribosome reaches the last codon in the AP, and again little force is exerted on the AP (*f*_*FL*_ ≈ 0). (B) Force profiles for the I27 domain (solid squares) and the non-folding (nf) mutant I27[W34E] (open squares). The standard error of *f*_*FL*_ is calculated for values of *L* where three or more experiments were performed.

### Cryo-EM shows that I27 folds in the mouth of the exit tunnel

To confirm that the peak in the force profile corresponds to the formation of a folded I27 domain, we replaced the SecM AP with the stronger TnaC AP (32–34) and purified stalled ribosome-nascent chain complexes (RNCs) carrying an N-terminally His-tagged I27[*L*=35] construct (see Methods). The construct was expressed in *E. coli*, RNCs were purified using the N-terminal His-tag, and an RNC structure with an average resolution of 3.2 Å (SI Appendix, Fig. S1) was obtained by cryo-EM. In addition to the density corresponding to the TnaC AP, a well-defined globular density (~4.5-9 Å resolution) was visible protruding from the exit tunnel (Figure 2A). Given the flat ellipsoidal shapes of the protruding density and of the I27 structure, there is only one way to fit the NMR structure of I27 (PDB 1TIT (35)) that gives a good Fourier-shell correlation between the isolated I27 density and the map generated from the I27 PDB model (SI Appendix, Fig. S2). In the fitted model, the C-terminal end of I27 extends into the exit tunnel and a β-hairpin loop on ribosomal protein uL24 is lodged in a cavity in I27 (Figure 2B and Supporting Video S1). The I27 domain further packs against ribosomal protein uL29 and ribosomal 23S RNA (Figure 2C), as if it is being pulled tight against the ribosome by the nascent chain. We conclude that the peak at *L* = 35-38 residues in the force profile indeed represents the cotranslational folding of the I27 domain at the tunnel exit.

**Figure 2.**
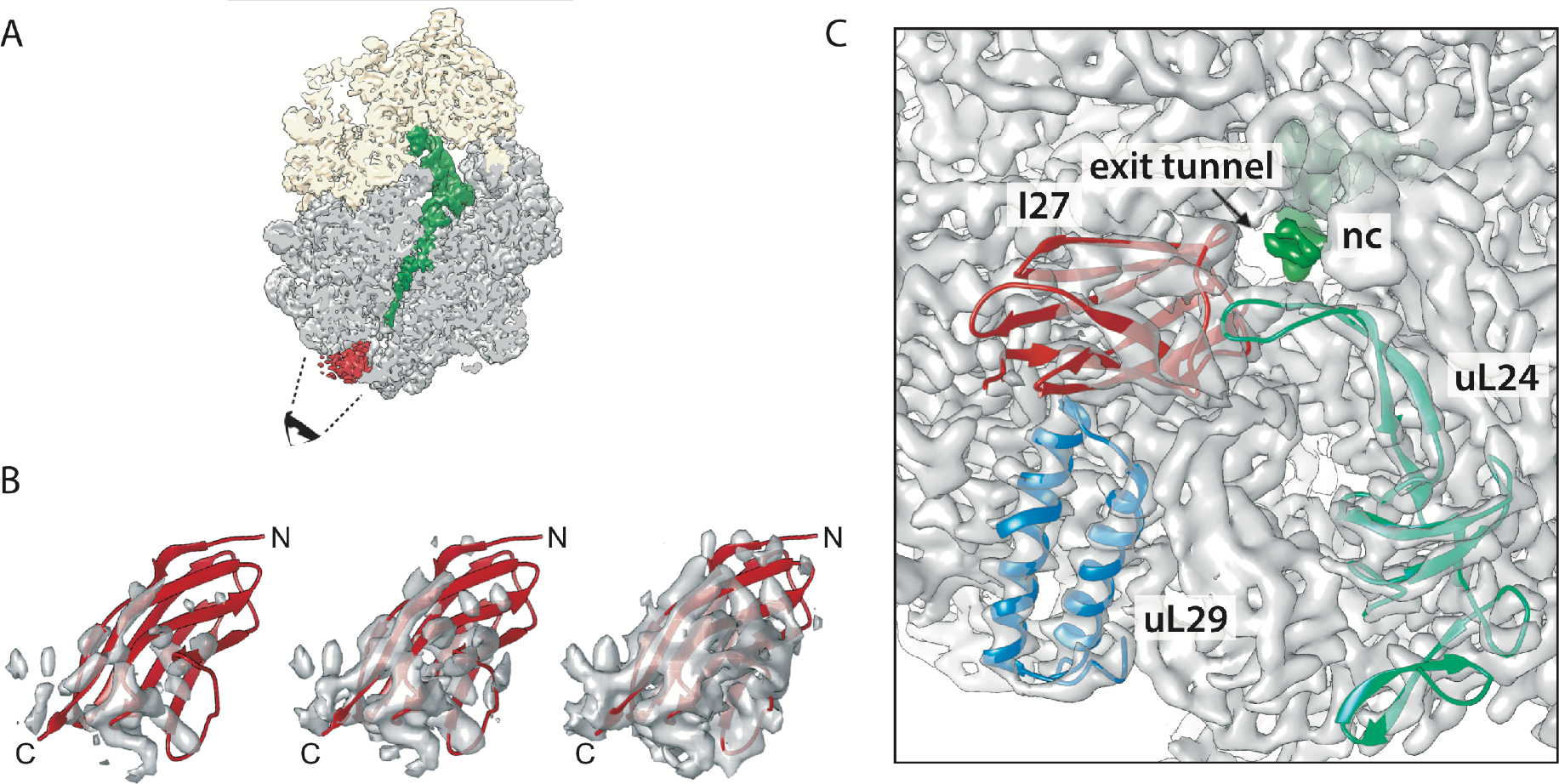
Cryo-EM structure of I27[*L*=35] RNCs. (A) Cryo-EM reconstruction of the I27-TnaC[*L* = 35] RNC. The ribosomal small subunit is shown in yellow, the large subunit in grey, the peptidyl-tRNA with the nascent chain in green, and an additional density corresponding to I27 at the ribosome tunnel exit in red. The black cartoon eye and dash lines indicate the angle of view in panel (C). The density contour level for feature visualization is at 1.7 times root-mean-square deviation (1.7 RMSD). (B) Rigid-body fit of the I27 domain (PDB 1TIT) to the cryo-EM density map displaying from high (left) to low (right) contour levels at 2.6, 2.0 and 1.4 RMSD, respectively. N and C represent the N and C termini of the I27 domain, respectively. (C) View looking into the exit tunnel (arrow) with density for the nascent chain (nc) in dark green. Ribosomal proteins uL29 (blue; PDB 4UY8), uL24 (light green; the β hairpin close to I27 domain was re-modelled based on PDB 5NWY) and the fitted I27 domain (red) are shown in cartoon mode; 23S RNA and proteins not contacting I27 are shown as density only. The density contour level is at 5 RMSD excluding tRNA, nascent chain and I27 domain, which are displayed at 1.7 RMSD.

### Coarse-grained molecular dynamics simulations recapitulate I27 folding on the ribosome

The yield of folded protein in arrest peptide experiments has been used as a proxy for the pulling forces that are exerted on the nascent chain at different points during translation in all studies to date (1, 2, 29). Here, to further elucidate the molecular origins of these forces and provide a quantitative interpretation of the observed folding yield of I27, we have calculated force profiles based on coarse-grained MD simulations (see Methods). Briefly, in the MD model, the 50S subunit of the *E. coli* ribosome (36) (PDB 3OFR) and the nascent chain are explicitly represented using one bead at the position of the Cα atom per amino acid, and three beads (for P, C4’, N3) per RNA base (Figure 3A). The interactions within the protein were given by a standard structure-based model (37–39), which allowed it to fold and unfold. Interactions between the protein and ribosome beads were purely repulsive (40) and the ribosome beads were fixed in space, as in previous simulation studies (18). I27 was covalently attached to unstructured linkers having the same sequences as those used in the force-profile experiments (Figure 3B) and the C terminus of the linker was tethered to the last P atom in the A-site tRNA (41) with a harmonic potential, allowing the force exerted by the folding protein to be directly measured. The potential chosen was stiff enough that displacements caused by typical pulling forces were smaller than 1 Å. For each linker length *L*, we used umbrella sampling to determine the average force exerted on the AP by the protein in the folded and unfolded states while arrested, as well as the populations of those two states (Figure 3C). We also estimated the folding and unfolding rates directly from folding/unfolding simulations.

**Figure 3.**
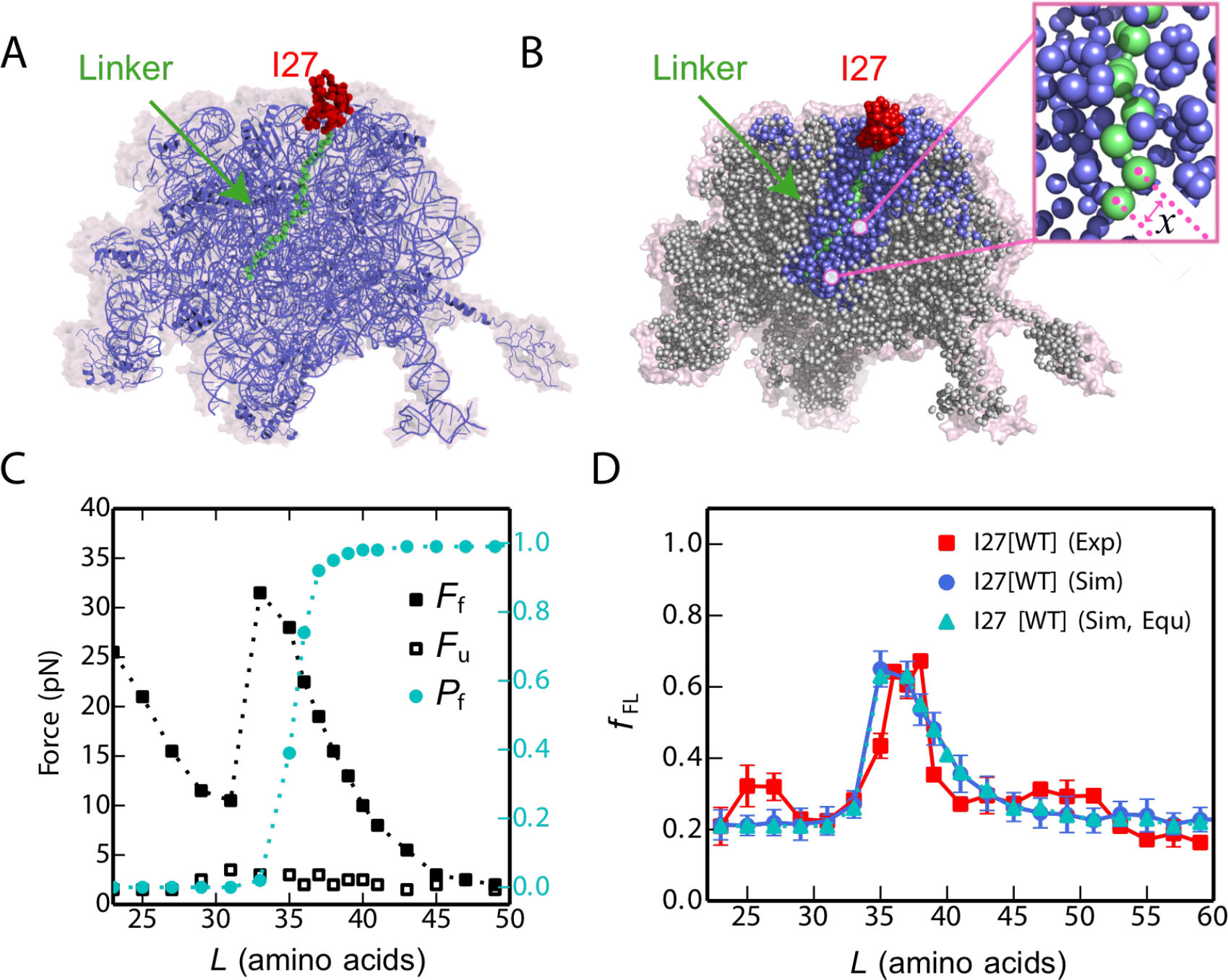
MD simulations of cotranslational folding of I27. (A) 50S subunit of the *E. coli* ribosome (PDB 3OFR) with I27[*L*=35] attached via an unstructured linker. (B) Coarsegrained model for I27 (red) and linker (green), with surrounding ribosomal pseudo-atoms in blue. Pseudo-atoms with grey colour are not used in the simulations. The instantaneous force exerted on the AP is calculated from the variation in the distance x between the C-terminal Pro pseudo-atom and the next pseudo-atom in the linker (see inset). (C) Average forces exerted on the AP by the unfolded state (*F*_*u*_, empty symbols) and folded state (*F*_f_, filled symbols) of I27 at different linker lengths *L*. The average fraction folded I27 for different *L*, *P*_f_, is shown in cyan on the right axis. Free energy profiles at each linker length are shown in SI Appendix, Fig. S4. (D) Experimental (red square) force profiles for cotranslational folding of I27. Force profiles calculated from simulations using full kinetic scheme or preequilibrium model are shown in blue circle and cyan triangle respectively. The RMSD of the *f*_*FL*_ between experiment and simulation is 0.08.

Given the experimentally-determined force-dependence of the escape rate *k*(*F*) (25), here approximated by a Bell-like model (42), we can calculate the expected escape rate while the protein is in the unfolded or folded state, from which the fraction full-length protein obtained with a given linker length and incubation time can be determined from a kinetic model, as described in Methods. The calculated *f*_*FL*_ profile for I27 is shown in Figure 3D (see also SI Appendix, Fig. S5) for the full solution of the kinetic model, as well as for an approximation in which the folding and unfolding rates are assumed to be faster than the escape rate (“preequilibrium”). Both results are very consistent with each other, as well as with the experimental profile. The peak in the folding yield arises as consequence of two opposing effects, the force exerted by the folding protein and population of the folded state, which respectively decrease and increase as the linker length increases. In the simulations with the I27[*L*=35] construct, the folded I27 domain is seen to occupy positions that largely overlap with the cryo-EM structure (Supporting Video S2). Overall, these results suggest that the MD model provides a good representation of the folding behaviour of the I27 domain in the ribosome exit tunnel. To show that the simulation model is not specific to I27, we have also applied it to another two proteins with different topologies for which experimental force profiles have been recorded, Spectrin R16 (all-α fold) and S6 (α/β fold) (2, 29). In these cases, we also obtain force profiles similar to experiment (SI Appendix, Fig. S6 and S7).

### Force profiles of I27 variants probe the folding pathway

To test whether the cotranslational folding pathway is the same as that observed for the isolated I27 domain *in vitro*, we investigated three destabilised variants of I27, both by simulation and experiment. One mutation in the core, Leu 58 to Ala (L58A), located in β-strand E (Figure 4A) destabilizes the protein by 3.2 kcal mol^−1^, and removes interactions that form early during folding of the isolated domain, playing a key role in formation of the folding nucleus (ϕ-value = 0.8) (26). Two further mutations, M67A and deletion of the N-terminal A-strand, remove interactions that form late in the folding of I27 (*i.e.*, both mutants have low ϕ-values (26, 27)). The A-strand is the first part of I27 to emerge from the ribosome, while M67 is located in a part of I27 that is shown by cryo-EM to be located in very close proximity to a β hairpin loop of ribosomal protein uL24 in I27-TnaC[*L*=35] RNCs (SI Appendix Fig. S3A). The interaction with the I27 domain shifts the tip of this uL24 hairpin by about 6 Å compared to its location in other RNC structures (SI Appendix Fig. S3B).

**Figure 4.**
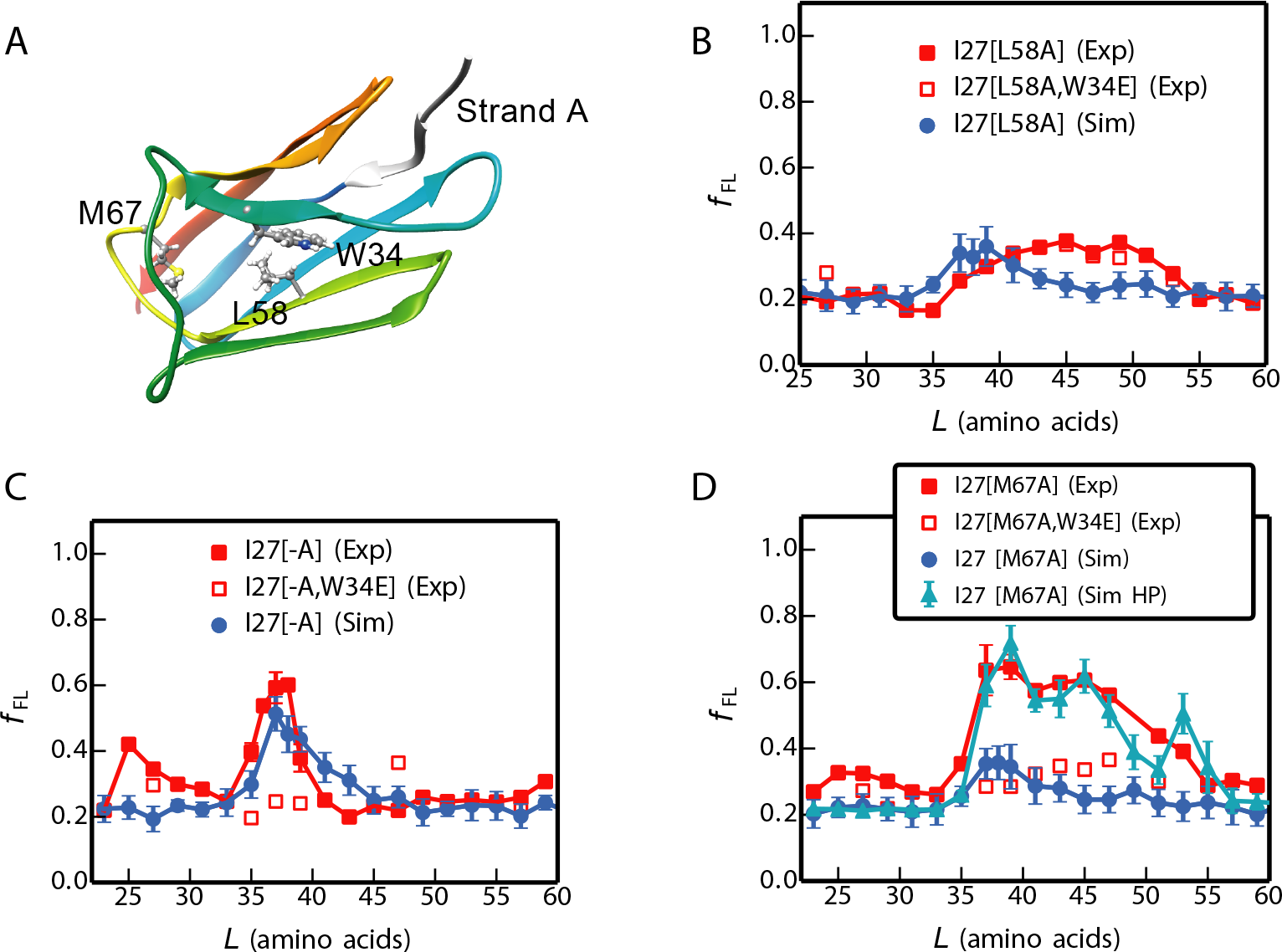
Simulations capture the experimental force profiles for mutant I27 domains. (A) Mutated residues in I27 (sticks). (B-D) Experimental (red) force profiles and calculated ones from full kinetic scheme (blue) for (B) I27[L58A], (c) A-strand deletion mutant I27[-A], (D) I27[M67A]. I27[M67A] (Sim HP) represents a simulation in which hydrophobic interactions between I27[M67A] and ribosome proteins uL23/uL29 are included. Experimental force profiles for non-folding mutants that contain an additional W34E mutation are shown as red open squares. The RMSD of the *f*_*FL*_ between experiment and simulation for I27[L58A], I27[−A] are 0.07 and 0.08 respectively. For I27[M67A], the *f*_*FL*_ RMSD is 0.07 between experiment and simulation (Sim HP).

The simulated force profile for the L58A variant predicts a much lower force peak than for wild-type I27; likewise, the experimental force peak is lower and broader than for wild-type, extending from *L* = 37-53 residues (Figure 4B). The fk values are very similar to those obtained for I27[L58A,W34E], a non-folding variant of I27[L58A]. Therefore, the weak forces seen at *L* ≈ 40-50 residues are not due to a folding event, indicating that I27[L58A] does not exert an appreciable force due to folding near the ribosome.

The A-strand comprises the first seven residues of I27 and removal of this strand, I27[-A], results in a destabilisation of 2.78 kcal mol^−1^; however, both the simulated and experimental force profiles for I27[−A] are very similar to those for wild-type I27 (Figure 4C). Residue M67 is located in the E-F loop, and mutation to alanine results in a destabilisation of 2.75 kcal mol^−1^; for this variant, folding commences at *L* ≈ 35 residues as for wild-type I27, but the peak is much broader (Figure 4D). Non-folding control experiments for variants I27[-A,W34E], and I27[M67A,W34E] (Figure 4C and D) show that the peaks in the force profiles for these variants are due to a folding event. These results show that deletion of the A-strand and destabilisation of the E-F loop do not affect the onset of cotranslational folding of I27, but that the M67A mutation increases the width of the folding transition. The simulation model used for the other mutants does not predict such a broad peak, suggesting that it may be necessary to include additional factors to reproduce the data for M67A. One possibility which may explain the result would be favourable interactions of the folded M67A mutant with the ribosome surface. The ribosomal surface proteins uL23 and uL29 have been suggested to form a potential interaction site for nascent proteins such as trigger factor (43), signal recognition particle (44) and SecYE (45). Here we have explored the hypothesis that the broad force peak of mutant M67A might due to interactions between an exposed hydrophobic cavity on I27[M67A] resulting from the mutation, and hydrophobic surface residues of ribosomal proteins uL23 and uL29. By introducing such interactions into the model, we are able to obtain a broad peak in the force profile very similar to that seen in experiment (Figure 4D).

### The folding pathway is only subtly affected by the presence of the ribosome

To compare the folding pathways when the protein is folding near the tunnel exit or outside the ribosome, we estimated ϕ-values based on the transition paths of I27 folding on the ribosome from our coarse-grained simulations, using a method introduced previously(46). The transition paths are those regions of the trajectory where the protein crosses the folding barrier, here defined as crossing between *Q* = 0.3 and *Q* = 0.7. For each linker length, 30 transition paths were collected from MD simulations. To reduce the uncertainty in the experimental reference data, we only compared with experimental ϕ-values if the change in folding stability between the mutant and the wild type is sufficiently large (|ΔΔG| > 7 kJ/mol) (47). As seen in Figure 5A, when the linker is long (L = 51 residues) and I27 is allowed to fold outside the ribosome, the calculated ϕ-values are consistent (Spearman correlation *r*=0.80) with the experimental values obtained for the folding of isolated I27 *in vitro* (26). For shorter linker lengths (*L* = 31 and 35 residues), calculated ϕ-values remain largely unchanged except for a slight increase near the N terminus (around residues 3-6) and a slight decrease near the C terminus (around residues 72-74) (Figure 5B and C).

**Figure 5.**
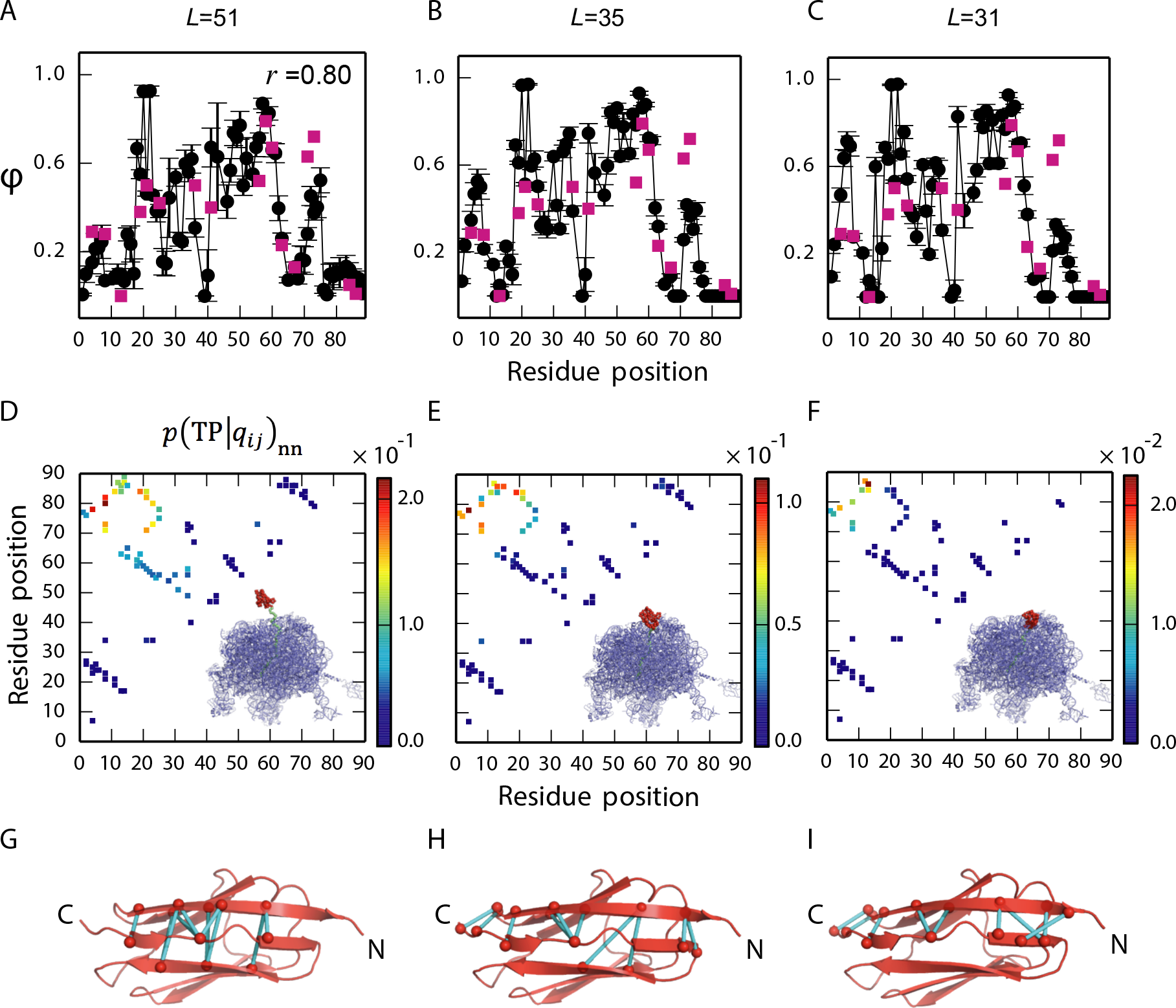
Simulated folding pathways for ribosome-tethered I27. LH column, *L*=51; middle column, *L*= 35; RH column, *L*=31. Top panels: Simulated ϕ-values for I27 (black). ϕ-values determined by *in vitro* folding of purified I27 are shown as red squares. At *L*=51 the simulated ϕ-values match well with experiment (spearman correlation *r*=0.80). At *L*=35 and *L*=31 the simulated ϕ-values are higher at the N terminus and lower at the C terminus, than the experimental values, reflecting a change in importance of these regions when I27 folds in the confines of the ribosome. Middle row: Relative probability that if a particular contact is formed then the protein is on a folding trajectory, *p*(TP|*q*_*ij*_)_nn_. When the protein is constrained the limiting factor is formation of a few key contacts. A cartoon of the ribosome with I27 in red is shown on each panel. Bottom row: The top ten most important contacts are coloured in cyan on the native structure.

To obtain a more detailed picture regarding the relative importance of different native contacts in the folding mechanism, we computed the conditional probability of being on a transition path (TP), given the formation of a contact *q*_*ij*_ between residues *i* and *j*, *p*(TP|*q*_*ij*_)_nn_ (48). This quantity indicates which native contacts are most important for determining a successful folding event. *p*(TP|*q*_*ij*_)_nn_ is closely related to the frequency of the contact qij on transition paths *p*(*q*_*ij*_|TP), but is effectively normalized by the probability that the contact is formed in non-native states *p*(*q*_*ij*_)_nn_ and can be expressed as:

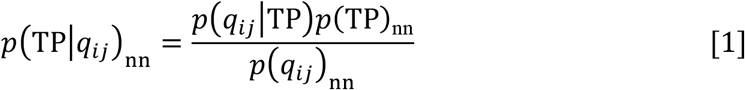

where *p*(TP)_nn_ is the fraction of non-native states which are on transition paths at equilibrium. The subscript nn means that only the non-native segments of a trajectory are included, *i.e.*, unfolded states and transition paths; the native, folded state is not included in the calculation since native contacts are always formed in this state. The simulations suggest that formation of native contacts between the N and C termini is somewhat more important when folding takes place in the mouth of the exit tunnel (*L* = 31 residues) than far outside the ribosome (*L* = 51 residues) (Figure 5D-F, upper left-hand corner in the panels). This is likely due to the greater difficulty of forming these contacts (examples are shown in Figure 5G-I) under ribosomal confinement; therefore, forming them becomes more critical in enabling the protein to fold.

## Discussion

Using a combination of MD simulation, force-profile measurements and cryo-EM, we have investigated the cotranslational folding pathway of the 89-residue titin I27 domain. I27 has been extensively characterised in previous *in vitro* folding studies (26, 27, 49–60). Results from all three techniques show that wild-type I27 folds in the mouth of the ribosome exit tunnel; in the cryo-EM structure of I27-TnaC[*L*=35] RNCs, I27 packs against ribosomal proteins uL24, uL29, and ribosomal 23S RNA. This is in apparent contrast to a previous NMR study on another Ig-like protein, in which the domain was shown to acquire its native fold (as reflected in the NMR spectrum) only when fully outside the ribosome tunnel, at *L* = 42-47 residues linker length (20).

In order to determine the molecular origin of the measured force profile, we performed molecular dynamics simulations of I27 folding on the ribosome, varying the length of the linker sequence between the arrest peptide and the I27 domain. We calculated the pulling force directly from the simulations and translated this into yield of folded protein using a kinetic model parameterized based on known release kinetics of the SecM AP. This enabled us to recapitulate the experimental arrest peptide force measurement profile, and therefore relate *f*_*FL*_ directly to the force exerted on the arrest peptide. Our simulations demonstrate the direct effect that the restoring force of the nascent chain can have on determining when the protein folds on the ribosome. We show that *f*_*FL*_ depends upon a combination of the force exerted by the folded protein and the fraction of folded protein at the given linker length *L*.

In order to relate how destabilization of regions that fold early and late in the isolated domain affects folding on the ribosome, we used simulations to predict the onset of folding in three mutant variants of I27. A previous ϕ-value analysis of I27 (26) showed that early packing of the structurally central β-strands drives the folding of this domain, while peripheral strands and loop regions pack later in the folding process. Mutations in the folding core (such as L58A) slow folding, whereas mutations in the periphery have no effect on folding rates (26). L58 is a key residue in the critical folding nucleus and almost fully packed in the transition state, in isolated domain studies. The simulated and experimental force profiles of I27 [L58A] show that this variant does not fold in or near the exit tunnel; hence, destabilisation of the central folding core prevents folding close to the ribosome. Since isolated I27[L58A] is fully folded, it is likely that this variant can only fold cotranslationally at longer linker lengths, when it is no longer in close proximity to the ribosome and exerts little force on the nascent chain.

Our experiments show that I27 variants destabilized in regions of the protein that are unstructured, or only partially structured, in the transition state, are still able to commence folding close to the ribosome. The force profiles reveal that the onset of folding of mutants with the A-strand deleted, or with the Met 67 to Ala mutation in the E-F loop, is the same as for wild-type although these have a similar destabilisation as L58A (Figure 4). The broader peak observed experimentally for M67A is harder to interpret. A plausible explanation is that the mutation introduces non-specific interactions of the folded domain with the ribosome surface, and we have shown that incorporating such interactions into the simulations could reproduce the results. An additional factor may be that that the mutation is in a region that interacts closely with ribosomal protein uL24 in the wild-type cryo-EM structure (SI Appendix Fig. S3).

Our simulations reproduce the onset of folding in the three mutant variants of I27 (Figure 4), and so give us the confidence to investigate how confinement within the ribosome affects the folding pathway of I27. We used simulations to investigate the folding of I27 arrested on the ribosome at various linker lengths, using a Bayesian method for testing the importance of specific contacts on the folding pathway, as well as by computing ϕ-values (Figure 5). Overall, we find that the mechanism and pathway of folding are robust towards variation in linker length and relatively insensitive to the presence of the ribosome; small but significant changes are observed only for contacts near the N and C termini. These shifts are consistent with the greater importance of forming N-terminal contacts when the C terminus is sequestered within the exit tunnel, possibly to compensate for loss of contacts at the C terminus.

In our kinetic modelling, we found that we obtained similar results with or without the assumption that folding and unfolding are fast relative to the escape rate, suggesting that this “pre-equilibrium” assumption is justified, at least for this protein. The reason for its validity in the case of I27 can be seen by comparing the folding and unfolding rates with the force-dependent escape rate of ~ 2.4 × 10^−3^ s^−1^ obtained at the highest forces of ~20 pN (c.f., Fig. 3C). Folding and unfolding rates at different linker lengths can be obtained by combining the linker-length dependence of the rates from simulation with the known folding/unfolding rates for isolated I27 from experiment (SI Appendix, Fig. S9). The presence of the ribosome increases the unfolding rate at shorter linker lengths so that it is faster than the maximum escape rate, while not slowing the folding rate sufficiently for it to drop below the escape rate. Note that the unfolding rate does drop below the maximum escape rate at larger linker lengths, but by that point the folded population is already almost 100%, so the preequilibrium assumption still gives accurate results. Although this assumption appears to be justified in the case of I27, it is probably not true in general, and it will be interesting to investigate for slower-folding proteins in future.

The arrest peptide experiments, in which a protein exerts a force due to folding in some ways resemble atomic force microscopy or optical tweezer experiments in which an external force is applied to the protein termini. It is important to note, however, that the nature and effect of the forces exerted on the folding protein by tethering to the ribosome are very different than is the case for pulling on both termini by an external force. For example, forces of the magnitude seen in this work (up to ~20 pN) tend to have very little effect on the unfolding rate when applied to the termini of I27, due to the similarity in extension of the folded and transition states (61); by contrast folding rates are dramatically slowed, even by very small forces, due to the large difference in extension of between unfolded and transition states (56). The forces arising from tethering to the ribosome are due to the folding of the protein itself rather than an external device. They arise from the constriction of available configuration space, particularly for folded and partially folded states, as well as from any attractive interactions between the protein and the ribosome. Our simulations suggest that for I27, reducing the linker length speeds up unfolding and slows folding rates by similar factors. Thus, it seems that comparisons to the effects of forces exerted by AFM and optical tweezer experiments need to be performed with care.

We have previously shown that α-helical proteins can fold co-translationally (2), perhaps unsurprising since helical structures are dominated by short-range interactions and helices can form within the ribosome tunnel itself (62, 63). Here, our equilibrium arrest-peptide assay and structural studies reveal that an all-β protein, titin I27, is able to fold within the mouth of the ribosome exit tunnel, despite its folding being dominated by long-range interactions. Molecular simulations, accounting for the effect of the entropic restoring force on protein stability, reproduce the yield of protein from experiments remarkably well. These simulations reveal that I27 folds on the ribosome by the same pathway as when the protein folds away from the confines of the ribosome. We note that a similar conclusion has been reached by Guinn et al. (64) for another small protein, src SH3, using a completely different experimental approach which combines optical tweezer experiments and chemical denaturant to characterize the folding pathway of src SH3. Thus, the evidence so far suggests that single-domain proteins, both α-helical and β-sheet, can fold close to the ribosome. On the other hand, while all-β proteins appear to fold by a similar pathway with or without the ribosome present, there is evidence for α-helical proteins forming partially structured cotranslational intermediates (11, 65) or folding by different pathways on the ribosome (2). This mechanistic difference may relate partly to the small contact order of helical proteins, allowing partially folded states to be more stable than for all-β proteins. The situation for multidomain proteins is likely to be still more complicated, as some studies have already indicated (11, 23, 66, 67).

## Acknowledgements

This work was supported by grants from the Knut and Alice Wallenberg Foundation, the Swedish Cancer Foundation, and the Swedish Research Council to GvH, by grants from the Deutsche Forschungsgemeinschaft (DFG) GRK 1721 and FOR1805 to RB, by a DFG fellowship through the Graduate School of Quantitative Biosciences Munich (QBM) to TS, and by the Wellcome Trust (WT095195) to JC; PT and RBB were supported by the Intramural Research Program of the National Institute of Diabetes and Digestive and Kidney Diseases of the National Institutes of Health; JC is a Wellcome Trust Senior Research Fellow. The cryo-EM data were collected at the Swedish National Cryo-EM Facility funded by the Knut and Alice Wallenberg Foundation, the Family Erling Persson Foundation and the Science for Life Laboratory. This work utilized the computational resources of the NIH HPC Biowulf cluster. (http://hpc.nih.gov)

## Materials and Methods

### Enzymes and chemicals

All enzymes were obtained from Thermo Scientific. Oligonucleotides were purchased from Life Technologies. In-Fusion Cloning kits were obtained from Clontech and DNA purification kits were purchased from Qiagen. PUREfrex cell-free translation system was obtained from Eurogentec. [35S]-methionine was purchased from Perkin Elmer. Instant Blue protein stain was purchased from Expedeon.

### DNA manipulation

Titin I27 constructs for *in vitro* translation were generated in pRSET A plasmid (Invitrogen) (previously modified to remove the sequence including the entire T7 gene 10 leader and EK recognition site up to, but not including, the *BamH* I site and replaced with a sequence encoding residues L, V, P, R, G, S) carrying the *E. coli* SecM arrest peptide (FSTPVWISQAQGIRAGP) and a truncated *E. coli lepB* gene, under the control of a T7 promoter. Increasing linker lengths were generated in pRSET A by PCR; linear pRSET A constructs (containing the SecM AP and truncated lepB, but lacking I27) were generated by PCR using primers which extended the linker from 23 aa to 63 aa (in steps of 2 aa) from the direction of the C to the N terminus. I27 flanked by GSGS linkers was amplified by PCR with overhanging homology to the plasmid containing the desired linker length. Cloning was performed using the In-Fusion system (Takara Bio USA, Inc.), according to the manufacturer’s instructions. The final two C-terminal residues (EL) of the 89 aa Titin I27 construct are not structured in the PDB file 1TIT, and are therefore included in the linker region. The amino acid sequence of the construct I27[*L*=63] is as follows (I27 in bold and SecM AP underlined): MRGSHHHHHHGLVPRGSGS**LIEVEKPLYGVEVFVGETAHFEIELSEPDVHGQWKLKGQPLAASPDCEIIEDGKKHILILHNCQLGMTGEVSFQAANTKSAANLKVK**EL SGSGKFAYGIKDPIYQKTLVPGQQNATWIVPPGQYFMMGDWMSS<u>FSTPVWISQAQGIRAGP</u>GS SDKQEGEWPTGLRLSRIGGIH**

The mutants I27[-A] (lacking β-strand A), I27[L58A] and I27[M67A] were generated for each linker length by site-directed mutagenesis. For the wild-type I27 and I27[-A] constructs with *L* = 27, 35, 37, 39, 47 and 57 residues, site-directed mutagenesis was performed to generate constructs with the non-functional FSTPVWISQAQGIRAG<u>A</u> arrest peptide (mutated residue underlined) as full-length controls, and constructs with the crucial Pro, at the end of the AP, substituted with a stop codon as arrest controls. Site-directed mutagenesis was performed to generate W34E variants as non-folding (nf) controls at *L* = 27, 29, 31, 35, 37, 39, 41, 43, 47, 49, 51 and 57 for wild-type I27; *L* = 27, 35, 37, 39, 47 and 57 residues for I27[-A]; *L* = 27, 41, 45, 47, 49 and 53 residues for I27[L58A]; *L* = 27, 29, 37, 39, 41, 43, 45, 47 and 51 residues for I27[M67A]. All constructs were verified by DNA sequencing.

### In vitro transcription and translation

Transcription and translation were performed using the commercially available PUREfrex *in vitro* system (GeneFrontier Corporation), according to the manufacturer’s protocol, using 250 µg plasmid DNA as template. Synthesis of [^35^S]-Met-labeled polypeptides was performed at 37 °C, 500 r.p.m. for exactly 15 min. The reaction was quenched by the addition of an equal volume of 10% ice-cold trichloroacetic acid (TCA). The samples were incubated on ice for 30 min and centrifuged for 5 min at 20,800 × *g* and 4 °C. Pellets were dissolved in sample buffer and treated with RNase A (400 μg ml^−1^) for 15 min at 37 °C before the samples were resolved by SDS-PAGE and imaged on a Typhoon Trio or Typhoon 9000 phosphorimager (GE Healthcare). Bands were quantified using ImageJ to obtain an intensity cross section, (http://rsb.info.nih.gov/ij/), which was subsequently fit to a Gaussian distribution using inhouse software (Kaleidagraph, Synergy Software). The fraction full-length protein, *f*_*FL*_ was calculated as *f*_*FL*_ = *I*_*FL*_(*I*_*FL*_+*I*_*A*_), where *I*_*FL*_ and *I*_*A*_ are the intensities of the bands representing the full-length and arrested forms of the protein. For wild-type I27 and six nf control samples (*L* = 27, 35, 37, 39, 47 and 57 residues), *in vitro* transcription and translation were also performed at 37 °C, 500 r.p.m. for exactly 30 min. The resultant force profile was slightly higher than that obtained at 15 min but has essentially the same shape (SI Fig. S5).

The reproducibility of force profile data has been discussed previously (2). For wild-type I27, data points *L* = 61 and 63 residues are a single experiment; *L* = 33, 36, 38, 45, 53, 55 and 59 residues are an average of 2 experiments; all other values of *L* are an average of at least 3 experiments. For I27[-A] strand, *L* = 23, 25, 33, 41, 43, 51, 53, 55 residues are a single experiment; all other values of *L* are an average of 2 experiments, except *L* = 35, 37 and 39 residues which are an average of at least 3 experiments. For I27[L58A], all data points are a single experiment except *L* = 27, 37, 41, 45, 47, 49 and 53 residues, which are an average of 2 experiments. For I27[M67A], *L* = 23, 25 and 51 - 63 residues are a single experiment; *L* = 29 - 35 residues are an average of 2 experiments; *L* = 27, 37 - 47 and 51 residues are an average of at least 3 experiments. For wild-type I27 samples incubated for 30 min, all data points are a single experiment except *L* = 27, 35, 37, 39, 47 and 57 residues, which are an average of 2 experiments. For non-folding controls, all data points are a single experiment except for wild-type I27 *L* = 29, 31, 39, 43 and 47 residues which are an average of 2 experiments.

### Cloning and purification of ribosome-nascent chain complexes

The I27 construct at *L* = 35, which is at the peak of *f*_*FL*_ (Figure 1B), was studied by cryo-EM. The SecM AP in these constructs was substituted with the TnaC AP (34) for more stable arrest, and the constructs were engineered to maintain a linker length of 35 amino acid residues. An N-terminal 8X His tag was introduced to enable purification. The amino acid sequence of the construct used was (I27 in bold and TnaC AP underlined): MDMGHHHHHHHHDYDIPTTLEVLF QGPGT**LIEVEKPLY GVEVFVGETAHFEIELS EPDVHGQWKLKGQPLAASPDCEIIEDGKKHILILHNCQLGMTGEVSFQAANTKS AANLKVK**EL SGSGSGS GGP<u>NILHISVT SKWFNIDNKIVDHRP</u>** The construct was engineered into a pBAD expression vector, under the control of an arabinose-inducible promoter. The translation-initiation region was optimized as described in (68). The plasmid was transformed into the *E. coli* KC6 *ΔsmpB ΔssrA* strain. 4 colonies were picked and tested for expression of the RNCs at 37°C in Lysogeny broth (LB).

Large-scale purification of RNCs was carried out based on a protocol described in (34). Briefly, a single colony of the KC6 cells found to express the RNCs was picked and cultured in LB at 37°C to an A600 of 0.5. Expression was induced with 0.3% arabinose and was carried out for 1 hour. Thereafter, the cells were chilled on ice, harvested by centrifugation, and resuspended in Buffer A at pH 7.5 (50 mM HEPES-KOH, 250 mM KOAc, 2 mM Tryptophan, 0.1% DDM, 0.1% Complete protease inhibitor). Cell lysis was carried out by passing the cell suspension thrice through the Emulsifex (Avestin) at 8000 psi at 4°C. The lysate was cleared of cell debris by centrifugation at 30,000xg for 30 min in the JA25-50 rotor (Beckman Coulter). The supernatant obtained was loaded on a 750 mM sucrose cushion (in Buffer A) and centrifuged at 45, 000 × g for 24 hours in a Ti70 rotor (Beckman Coulter) to obtain a crude ribosomal pellet, which was resuspended in 200 μl Buffer A by shaking gently on ice.

RNCs from the crude suspension were purified via their His tags by affinity purification using Talon (Clontech) beads, which was pre-incubated with 10 μg/ml tRNA to reduce unspecific binding of ribosomes. The suspension was incubated with the beads for 1 hour at 4°C and subsequently washed with 20 column volumes of Buffer B at pH 7.5 (50 mM HEPES-KOH, 10 mM Mg(OAc)_2_, 0.1% Complete Protease Inhibitor, 250 mM sucrose, 2 mM Tryptophan). RNCs were eluted by incubating the Talon beads with Buffer C at pH 7.5 (50 mM HEPES, 150 mM KOAc, 10 mM Mg(OAc)_2_, 0.1% Complete protease inhibitor, 150 mM imidazole, 250 mM sucrose) for 15 minutes and subsequently collecting the flowthrough. Elution was carried out thrice and the eluents were concentrated by centrifugation at 40,000 rpm for 2.5 hours in a TLA 100.3 rotor (Beckman Coulter). The pellet obtained at the end of this step was gently suspended in a minimal volume of Buffer D at pH 7 (20 mM HEPES-KOH, 50 mM KOAc, 5 mM Mg(OAc)_2_, 125 mM sucrose, 2 mM Trp, 0.03% DDM).

### Cryo-EM sample preparation, data collection, processing and accession codes

Approximately 4 A_260_/ml units of RNCs were loaded on Quantifoil R2/2 grids coated with carbon (3 nm thick) and vitrified using the Vitrobot Mark IV (FEI-Thermo) following the manufacturer’s instructions. Cryo-EM data was collected at the Cryo-EM National Facility at the Science for Life Laboratory in Stockholm, Sweden.

Data was acquired on a 300 keV Titan Krios microscope (FEI) equipped with a K2 camera and a direct electron detector (both from Gatan). The camera was calibrated to achieve a pixel size of 1.06 Å at the specimen level. 30 frames were acquired with an electron dose 0. 926 e^−^/Å^2^/frame and a total dose of 27.767 e^−^/Å^2^ and defocus values between −1 to −3 µm. The first two frames were discarded and the rest were aligned using MotionCor2 (69). Raw images were cropped into squares by RELION 2.1 beta 1 (70). Power-spectra, defocus values and estimation of resolution were determined using the Gctf software (71) and all 2,613 micrographs were manually inspected in real space, in which 2,613 were retained. 468,015 particles were automatically picked by Gautomatch (http://www.mrc-lmb.cam.ac.uk/kzhang/) using the *E. coli* 70S ribosome as a template. Single particles were processed by RELION 2.1 beta 1 (70). After 80 rounds of 2D classification, 384,039 particles were subjected to 3D refinement using the *E. coli* 70S ribosome as reference structure, followed by 160 rounds of 3D classification without masking and 25 rounds of tRNA-focused sorting. One major class containing 301,510 particles (64% of the total) was further refined including using a 50S mask, resulting in a final reconstruction with an average resolution of 3.2 Å (0.143 FSC). The local resolution was calculated by ResMap (72). Finally, the final map was obtained by local B-factoring followed by low-pass filtering to 4.5 Å by RELION 2.1 beta 1 (70) in order to best demonstrate the I27 domain.

For interpretation of the cryo-EM density, the cryo-EM structure model (PDB 4YU8) of *E. coli* TnaC-stalled ribosome was fitted into corresponding density using UCSF Chimera (73). The NMR model (PDB 1TIT) of I27 domain was fitted into the extra density of TnaC-stalled ribosome using UCSF Chimera (73). Since the I27 domain represents a flat ellipsoid, we used all four major and minor axes covering all possible orientations of the model fitting within the density to validate the orientation of the fitted I27 model. Briefly, the model with four different orientations were converted into densities (8 Å) by UCSF Chimera, and the crosscorrelation coefficients of each model map and the isolated I27 density were calculated by RELION 2.1 beta 1 (70). Finally, uL24 β hairpin was remodeled as the tip of the hairpin is shifted due to the existence of I27 domain.

Figures showing electron densities and atomic models were generated using UCSF Chimera (73). Electron densities are shown at multiple contour levels in Figure 2 and SI Appendix, Fig. S1. The contour levels relative to the root-mean-square deviation (RMSD) were calculated from the final map values. Final map contains the volume for the entire RNC including the I27 domain.

Coordinates for the cryo-EM map of the ribosome with the I27 domain density have been deposited at the EMDataBank under accession code EMD-*xxxx*. Coordinates of fitted *E.coli* TnaC-stalled ribosome (PDB 4UY8; uL24 remodeled) and I27 domain (PDB 1TIT) models for interpreting the cryo-EM map have been deposited at the ProteinDataBank under accession code *xxxx*.

### Coarse-grained molecular simulations

The 50S subunit of the *E. coli* ribosome (PDB 3OFR (36)) and the nascent chain are explicitly represented using one bead at the position of the α-carbon atom of each amino acid, and three beads (for P, C4’, N3) per RNA residue (Figure 2A). The interactions within the protein were given by a standard structure-based model (37–39), which allowed it to fold and unfold. Interactions between the protein and ribosome beads were purely repulsive (40) and given by the same form of potential as for the structure-based model(37–39),

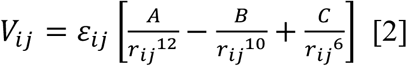

where *r*_*ij*_ is the distance between two beads *i* and *j*, ε_*ij*_ (=0.001 kJ/mol) sets the strength of the repulsive interactions. The amino acid, phosphate, sugar and base are assigned radii σ_*i*_ = 4.5, 3.2, 5.1 and 4.5 Å respectively, and coefficients in Eq. 2 for interactions between protein and ribosome beads *i,j* are obtained from the mixing rules 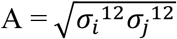, 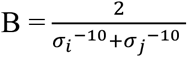 and 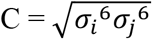.

During the simulations, the positions of the ribosome atoms were fixed in space, as in previous studies (18). The linker between the AP and I27 was tethered by its C terminus to the last P atom of the A-site tRNA, but was otherwise free to fluctuate. The trajectory was propagated via Langevin dynamics, with a friction coefficient of 0.1 ps^−1^ and a time step of 10 fs, at 291 K in a version of the Gromacs 4.0.5 simulation code, modified to implement the potential given by Eq. 2 (74). All bonds (except the one used to measure force, below) were constrained to their equilibrium length using the LINCS algorithm (75). The attractive interactions between I27[M67A] and the hydrophobic residues (A, V, L, I, F, M, Y, W) on the surface of uL23 and uL29 are modelled as (76):

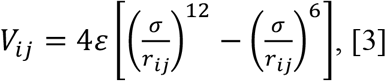

where *r*_*ij*_ is the distance between residues *i* and *j*, *σ* is the range of the interaction and ε; represent the strength of the interaction. *σ* and ε are fixed at 6 Å and 5 kJ/mol respectively. Residues of I27[M67A] which are involved in the attractive interactions are defined as the ones whose heavy atoms are within 4.5 Å of any heavy atoms from residue 67 in the native state.

To calculate the pulling force exerted on the nascent chain by the folding of I27, the bond between the last and the second last amino acid of the SecM AP was modelled by a harmonic potential as a function the distance between these two atoms, *x* (Figure 3B):

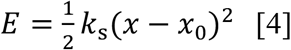

where *x*_0_ is a reference distance. Here *x*_0_ is set to 3.8 Å, which is the approximate distance between adjacent C_α_ atoms in protein structures and *k*_*s*_ is a spring constant, set to 3000 kJ.mol^−1^.nm^−2^. The value of *k*_*s*_ was chosen so that the average displacement *x* − *x*_0_ remains below 1 Å for forces up to ~500 pN, which is much larger than the forces actually exerted by the folding protein. The pulling force on the nascent chain was measured by the extension of this bond as *F* = −*k*_*s*_(*x* − *x*_0_).

I27 was covalently attached to unstructured linkers having the same sequences as used in the force-profile experiments (see Figure 2B). Linker amino acids are repulsive to both the ribosome and I27 beads, with interaction energy as described in Eq. 2.

The protein in its arrested state is subject to force *F*(*t*), which will fluctuate, for example when the protein folds or unfolds. The rate of escape from arrest has been shown to be force-dependent (25); here we approximate the sensitivity to force using the phenomenological expression originally proposed by Bell (42)

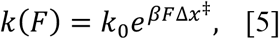

where *k*_0_ is a zero-force rupture rate, Δ*x*^‡^ is the distance from the free energy minimum to the transition state, *β* = 1/*k*_*B*_*T* where *k*_B_ is Boltzmann’s constant and *T* the absolute temperature. While there are functions to describe force-dependent rates with stronger theoretical basis, we use the Bell equation due to its simplicity and because its parameters have previously been estimated from optical tweezer experiments for the SecM AP. In all cases, we set *k*_0_ (Eq. 5) to 3.4 × 10^−4^ s^−1^ and Δ*x*^‡^ to 3.2 Å, based on the values determined by Goldman *et al.* (they estimated *k*_0_ and Δ*x*^‡^ to be in the range of 0.5 × 10^−4^ to 20 × 10^−4^ s^−1^ and 1-8 Å, respectively) (25).

We assume the probability of remaining on the ribosome *S*(*t*) = 1 − *f*_*FL*_(*t*) assuming that 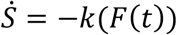 hence

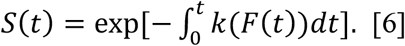

The escape of I27 from the ribosome can be described using kinetic model shown in SI Appendix, Fig. S8 which explicitly takes into account the linker length-dependent folding/unfolding rates of the I27 nascent chain, *k*_u_(*L*) and *k*_f_(*L*) on the ribosome, and the force-dependent rate of escape from ribosome: *k*(*F*_u_(*L*)) and *k*(*F*_f_(*L*)). To estimate *kf* at different linker lengths, we first carried out unbiased MD simulations to estimate the mean first passage time for folding 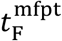, from which the folding rate can be calculated as 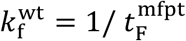. Similarly, the unfolding rate can be calculated from unfolding simulations as 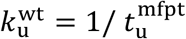. Since the rates in coarse-grained simulations are naturally much faster than in experiment, we globally scale the unfolding rates 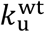 (*L*=21, 23 …61) at different linker lengths so that 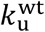 at very long linker lengths (*L*=61) is equal to the unfolding rate of isolated I27 (4.9×10^−4^ S^−1^). Similarly, 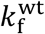(*L*=21, 23 …61) is scaled so that the 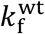 of I27 RNC[*L*=61] is equal to the unfolding rate of isolated I27 (SI Appendix, Fig. S9). For consistency with our pre-equilibrium solution, we further scale 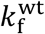 to match the stability of I27 RNC[*L*=61] in our simulation model, yielding 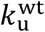 and 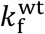 at *L*=61 of 4.9×10^−4^ S^−1^ and 0.14 S^−1^ (SI Appendix, Fig. S9) respectively. The same scaling method has been applied to the folding and unfolding rates of all mutants (I27[L58A], I27[M67A] and I27 [-A]) so that the folding/unfolding rates of the mutant RNC are consistent with the relative experimental values measured for the isolated mutants(26).

With the rates obtained from above, the time dependent survival probability *S*(*t*) is estimated by the kinetic Monte Carlo method (the Bortz-Kalos-Lebowitz algorithm (77)). The system is initialized at the state when the unfolded nascent chain just emerges from the ribosome tunnel (UA, SI Appendix, Fig. S8) at time t=0. At each Monte Carlo step, a uniform random number *δ* between 0 and 1 is chosen, and a transition from the current state state *s* to state *j* will occur for the state *j* which satisfies 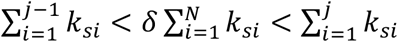, where *k*_*si*_ represents the transition rate from state *s* to state *i*. The time is updated by *t* = *t* + Δ*t*, where 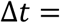 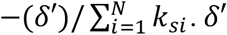 is a new number randomly chosen between 0 to 1.

The solution to the kinetic model can be simplified if we further assume that the escape from the ribosome is slow relative to the folding and unfolding of the protein. In this situation, we can approximate *S*(*t*) in terms of the mean forces experienced when the protein is unfolded, *F*_u_, or folded, *F*_f_, and the unfolded and folded populations of *P*_u_ and *P*_f_ respectively,

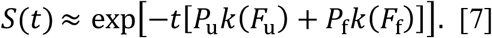

The equilibrium properties of the system for each linker length were obtained from umbrella sampling using the fraction of native contacts *Q* as the reaction coordinate, allowing *P*_u_, *P*_f_ and *F*_u_, *F*_f_ to be determined (Figure 3C). The details of the definition of *Q* have been previously described (48); in short, *Q* is defined as

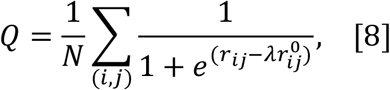

where the sum runs over the N pairs of native contacts (*i,j*), *r*_*ij*_ is the distance between *i* and *j* in configuration, 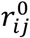 is the distance between *i* and *j* in the native state, λ =1.2 which accounts for fluctuations when the contact is formed. A boundary of *Q* = 0.5 is used to separate folded from unfolded states.

In order to characterize folding mechanism, we used transition paths from folding simulations for the *L* = 51 case at 291 K. 50 independent simulations, each started from fully extended configurations, were carried out for 4 microseconds. The folding barriers for the *L* = 31 and 35 cases are very high at the same temperature, therefore the transition paths are obtained from unfolding simulations instead. Starting from native-like folded configurations, 50 unfolding simulations were carried out, with each trajectory being 4 microseconds long. Transition paths were defined as those portions of the simulation trajectory from the last time I27 samples the configuration with *Q* < 0.3 till the first time it samples a configuration with Q > 0.7 (in the folding direction; opposite for unfolding). ε-Values were computed from the transition paths using the approximation:

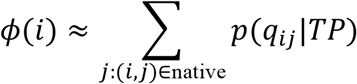

In which *p*(*q*_*ij*_|TP) is the probability that the native contact *q*_*ij*_ between residues *i* and *j* is formed on transition paths as defined above. We also characterized the importance of individual contacts in determining the folding mechanism using *p*(TP|*q*_*ij*_)_nn_ defined in Eq. 1 of the main text, i.e. the probability of being on a transition path given that contact *q*_*ij*_ is formed and the protein is not yet folded. Having already calculated *p*(TP|*q*_*ij*_) above, evaluating *p*(TP|*q*_*ij*_)_nn_ required *p*(*q*_*ij*_)_nn_, the probability of a contact being formed in all non-native fragments of the trajectory, and *p*(*TP*), the fraction of time spent on transition paths. For *L*=51, we obtained *p*(*q*_*ij*_)_nn_ directly from unbiased folding simulations, using the portion of the trajectory up to the first folding event (i.e. the first time *Q* > 0.7). For *L* =31 or 35, where the protein is still relatively unstable, we determined it from unfolding simulations by computing *p*(*q*_*ij*_) separately for the unfolded and transition-path portions of the trajectory and combining them weighted by *p*(TP)_nn_. We determined *p*(TP)_nn_ via folding (*L* = 51 case) and unfolding (*L* = 31 and *L* = 35 cases) simulations (described above). For the *L* = 51 case, 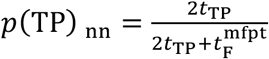 where *t*_TP_ is the mean transition path time and 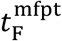 is the mean first passage time for folding obtained from the maximum likelihood estimator 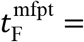 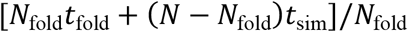 where *N* is the total number of trajectories (*N* = 50), *N*_fold_ is the number of trajectories folding within 4 μs, *t*_fold_ is the average folding time (of the trajectories which fold), and *t*_sim_ is the length of the simulations (4 μs). For the *L* = 31 and *L* = 35 cases, it is less efficient to obtain the folding time 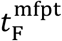 directly, therefore we estimate it based on the mean first passage time for unfolding, 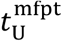, from unfolding simulations. 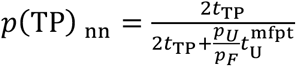 where *p*_u_ and *p*_f_ are the equilibrium populations of the unfolded and folded respectively determined from umbrella sampling.

## Supplementary Figures

**Figure S1.**
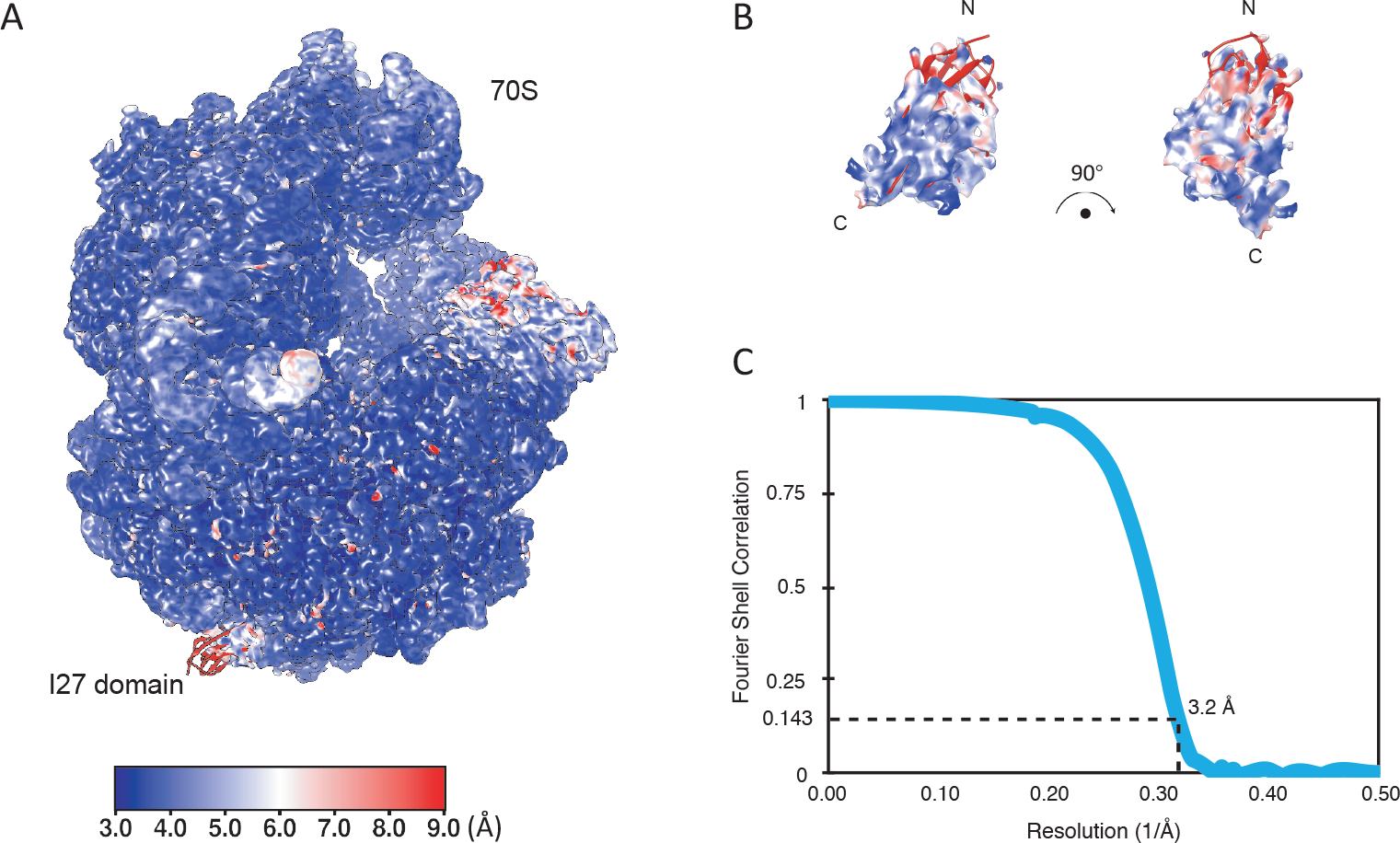
Resolution of the ribosome-nascent chain complex (RNC). (A) Calculation of the local resolution using Resmap (Kucukelbir, A. et al. Nat Methods 11, 63-65, 2014). The RNC density is displayed at 1.7 RMSD. (B) local resolution of the I27 domain. The I27 domain density is displayed at 2 RMSD. N and C termini are indicated. (C) Fourier-shell correlation (FSC) curve of the refined final map of the RNC, indicating the average resolution of 3.2 Å (at 0.143).

**Figure S2.**
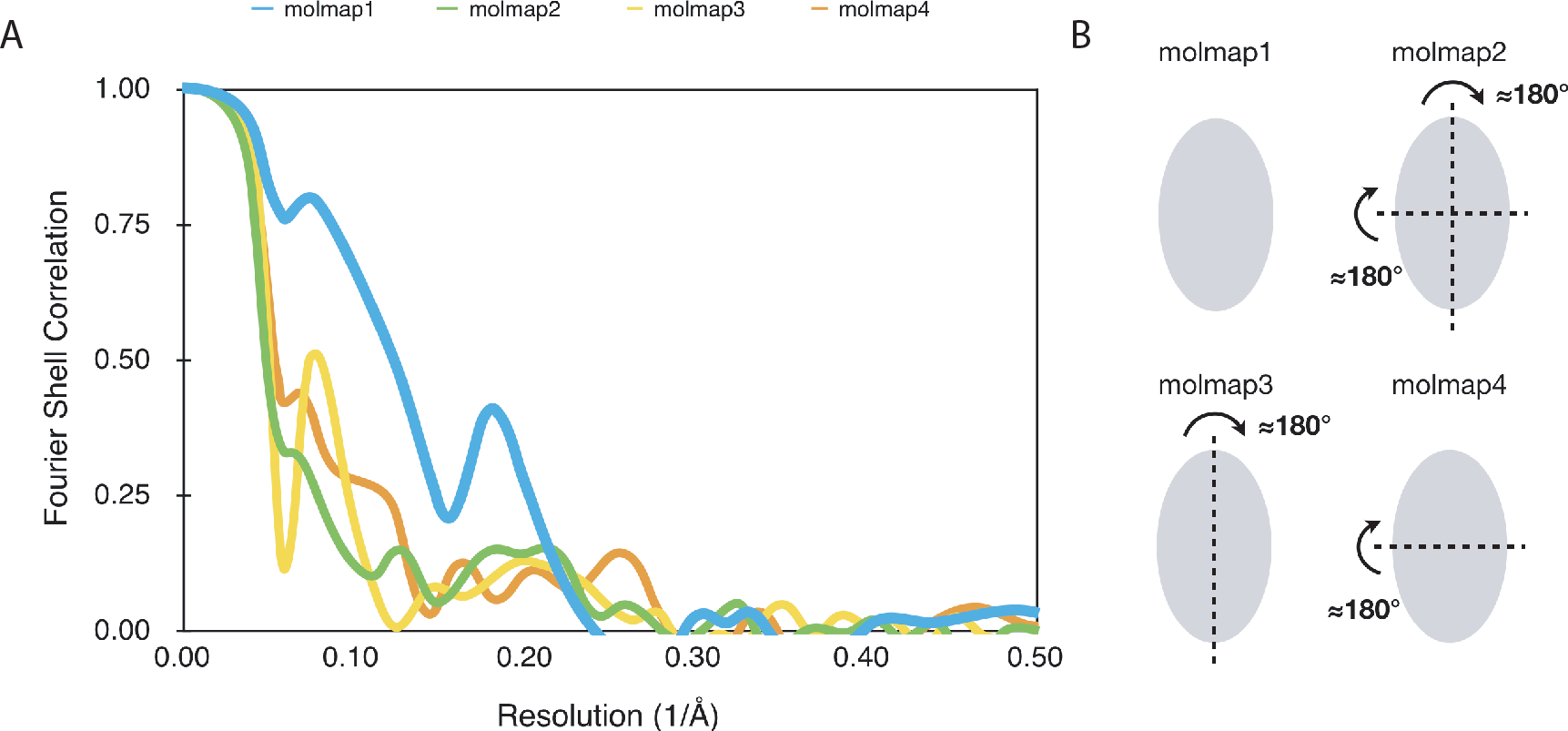
Validation of model orientation for I27 domain. To validate the orientation of the I27 domain model (PDB 1TIT) to its corresponding density, four possible orientations were tested. (A) The Fourier-shell correlations between the isolated I27 density and the map generated from the model of the final orientation (molmap1, blue) and the models fitted with the other three possibilities (molmap2, green; molmap3, yellow; molmap4, orange) were plotted. In the frequency range 0 to 0.2 (1000 to 5 Å) the correlation of molmapi is significantly higher compared to all other orientation molmaps. (B) The illustration showing the relationship among the four model orientations. Since the density represents a flat ellipsoid, we used all four major and minor axes covering all possible orientations of the model fitting within the density.

**Figure S3.**
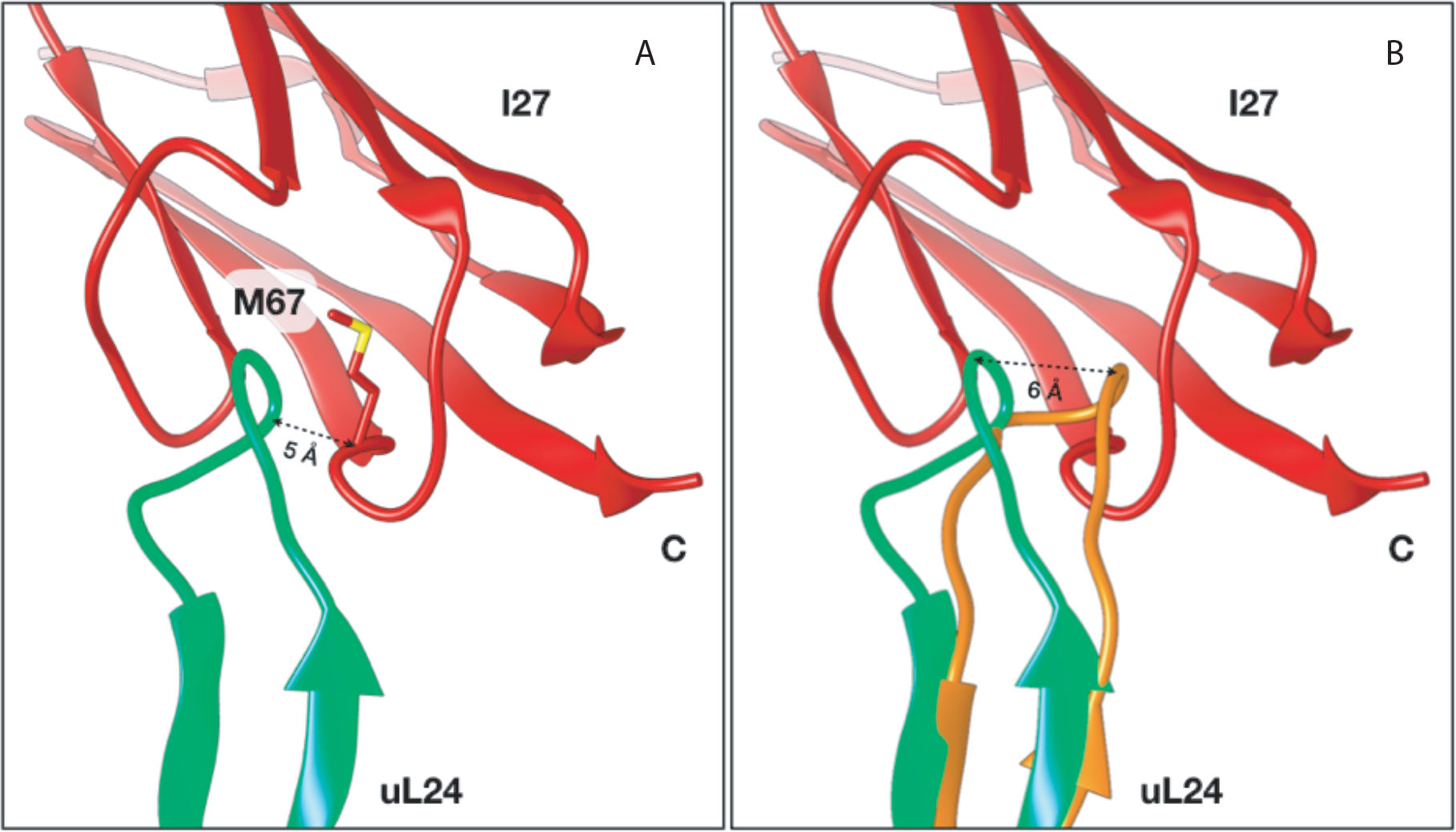
The I27 domain and a P hairpin in ribosomal protein uL24 close to the ribosomal exit tunnel. (A) Residue M67 in the I27 domain is located in close proximity to a β hairpin loop in uL24 in the cryo-EM structure of I27-TnaC[*L*=35] RNCs. (B) The uL24 β hairpin in the I27-RNC (light green; re-modeled based on PDB 5NWY) is ~ 6 Å shifted (distance measured via the backbone of Pro50) compared to its location in the VemP-RNC (orange; PDB 5NWY) and the TnaC-RNC (PDB 4UY8, not shown). C represents the C terminus of the I27 domain.

**Figure S4.**
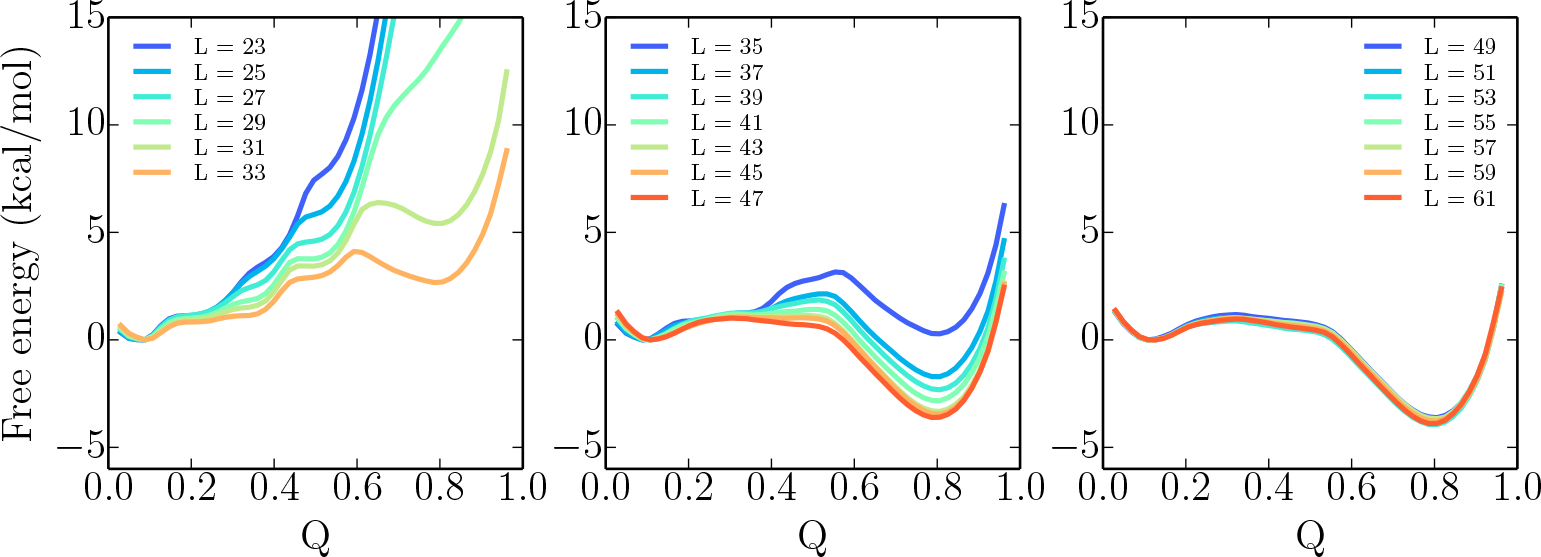
Simulation free energy *F*(*Q*) projected on the fraction of native contacts, *Q*, for I27 folding with different linker lengths (as indicated in legend) at 291K.

**Figure S5.**
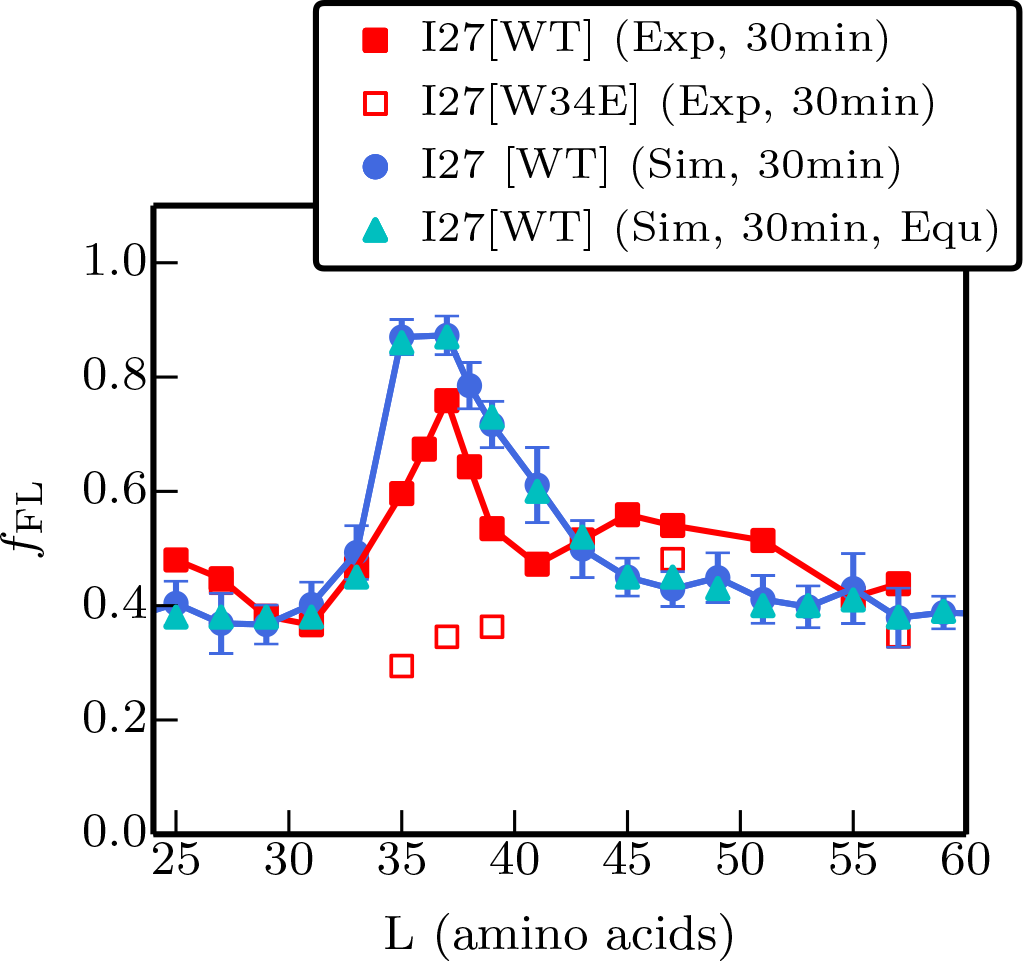
Experimental (red) and simulated profiles of fraction full length protein, *f*_*FL*_ obtained with a 30 min incubation. Note the higher background values compared to main text Figures 1 and 3D. Force profiles calculated 7rom simulations using full kinetic scheme and pre-equilibrium model are shown in blue circles and cyan triangles respectively. The RMSD of the *f*_*FL*_ between experiment and simulation is 0.12.

**Figure S6.**
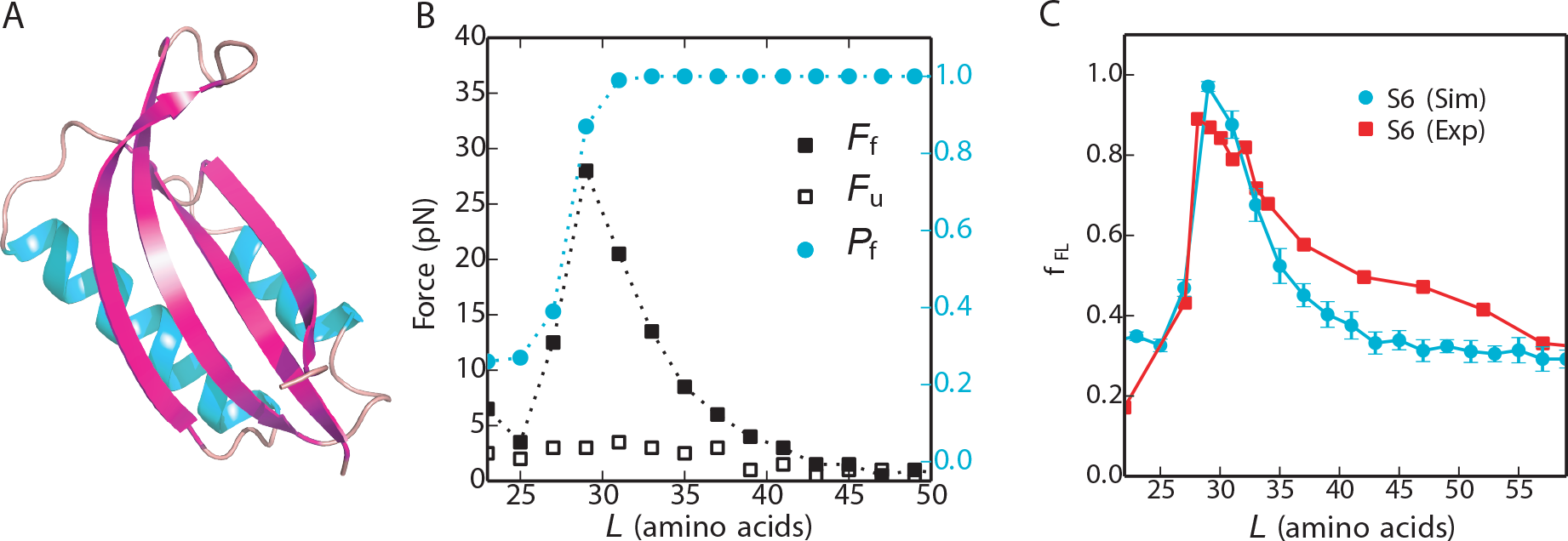
(A) Native structure of protein S6 (pdb code: 2KJV (78)) B) Average forces exerted on the AP by the unfolded state (F_f_, filled black symbols) and folded state (F_u_, empty black symbols) of S6 at different linker lengths *L*. The average fraction folded S6 for different *L*, P_f_, is shown in cyan on the right axis. (C) Experimental (red) and simulated (cyan) force profiles for cotranslational folding of S6 based on pre-equilibrium kinetic solution.

**Figure S7.**
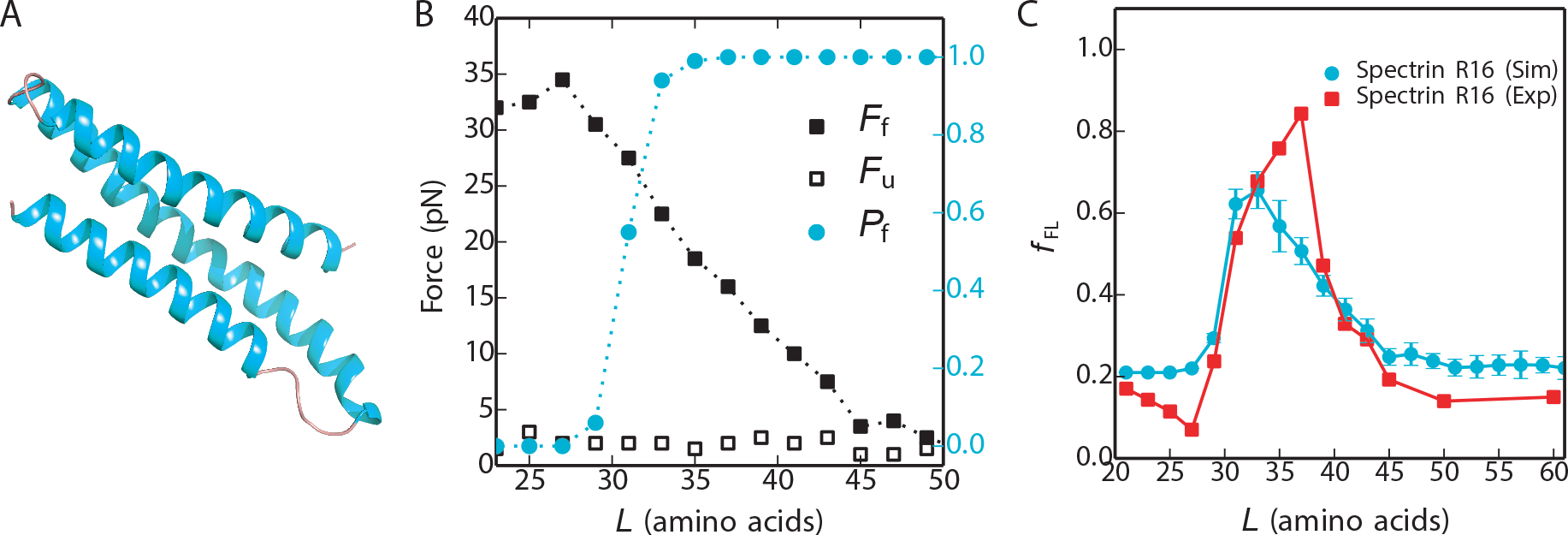
(A) Native structure of Spectrin R16 domain (PDB 1AJ3 (79)). (B) Average forces exerted on the AP by the unfolded state (F_f_, empty black symbols) and folded state (F_u_, filled black symbols) of R16 at different linker lengths *L*. The average fraction folded R16 for different *L*, P_f_, is shown in cyan on the right axis. (C) Experimental (red) and simulated (cyan) force profiles for cotranslational folding of R16 based on the pre-equilibrium kinetic solution.

**Figure S8.**
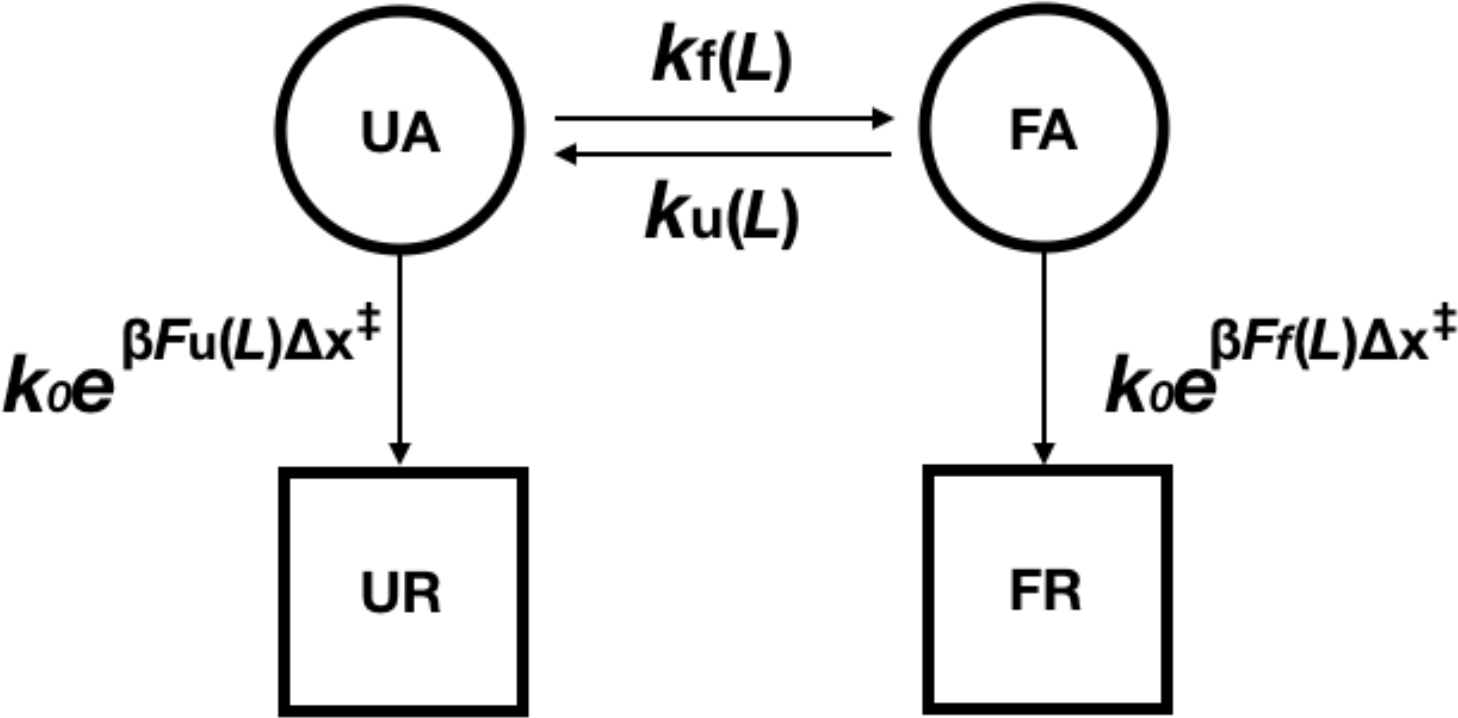
Schematic for the full kinetic model which describes the escape pathway of I27 from the ribosome. *k*_f_ and *k*_u_ are the linker length-dependent folding and unfolding rates respectively. UA: I27 is unfolded and arrested by ribosome. FA: I27 is folded and arrested by ribosome. UR: I27 is unfolded and released from ribosome. FR: I27 is folded and released from ribosome.

**Figure S9.**
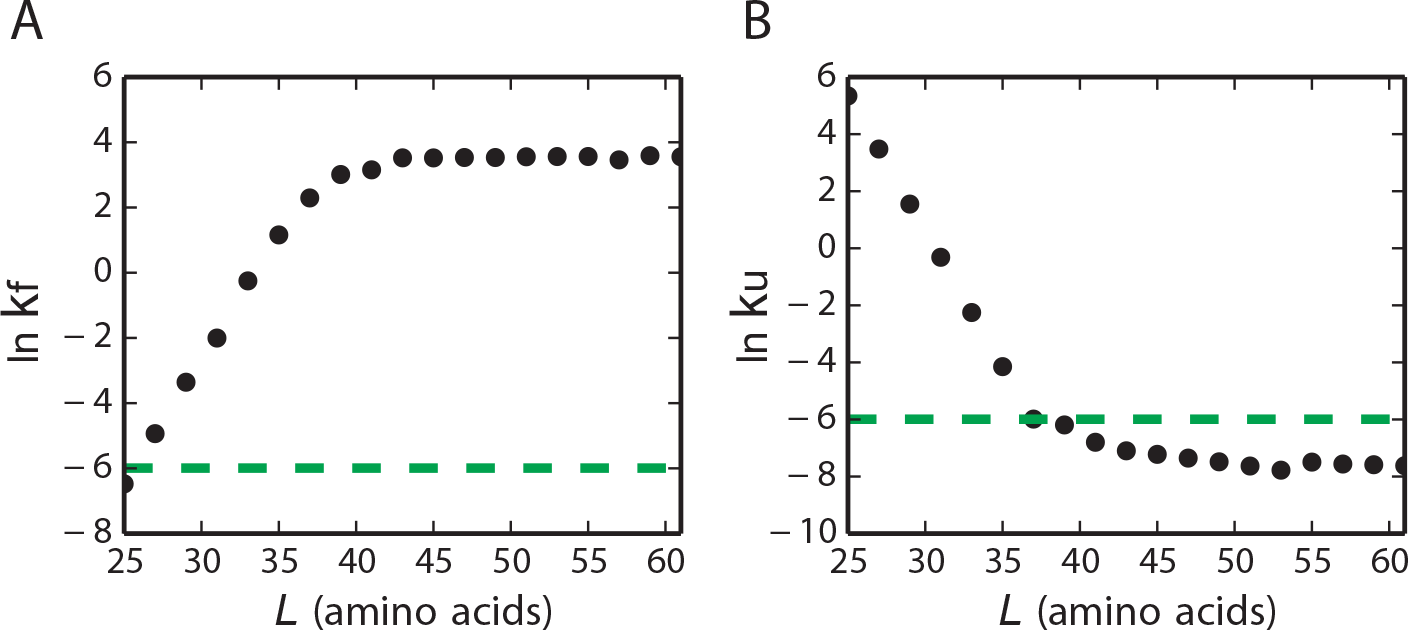
Dependence of folding rate *k*_f_ (left) and unfolding rate *k*_u_ (right) on the length of the linker between the AP and I27. Rates determined directly from simulations have been scaled so that *k*_f_ and *k*_u_ at large linker lengths are equal to the experimental values determined for the isolated protein. The green dashed line indicate the force-dependent escape rate of ~ 2.4 × 10^−3^S^−1^ obtained at the force of 20 pN.

**Supporting Video S1.**
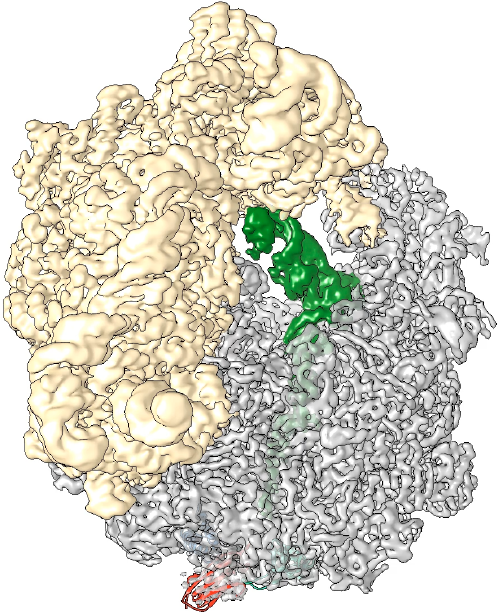
Cryo-EM density of ribosome and I27 (one static frame of the video). Video showing cryo-EM map for I27[*L*=35] RNCs. 30S in yellow, 50S and I27 domain in grey, tRNA and nascent chain in green, the model (PDB 1TIT) of I27 domain in red.

**Supporting Video S2.**
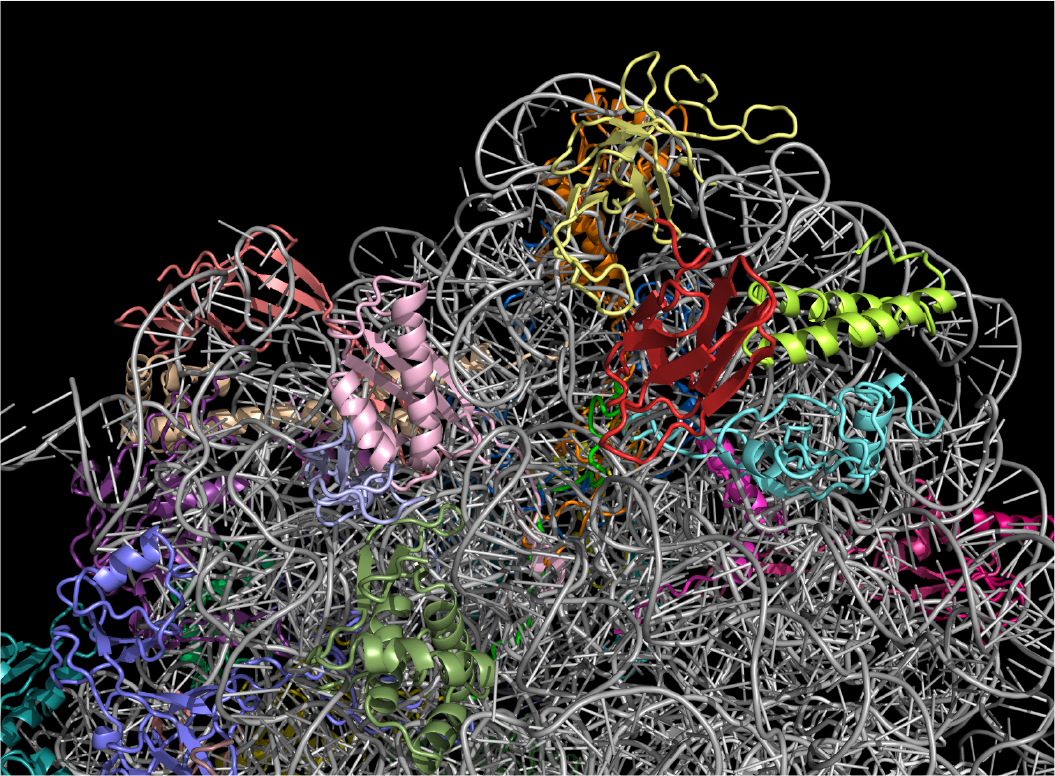
MD folding simulation (one static frame of the video). Video showing an unbiased 1.8 μsec fragment of an MD trajectory of I27 folding and unfolding at linker length *L*=35. Ribosomal 23 s rRNA is shown in white cartoon mode, ribosomal proteins uL24, uL29, uL23 are shown in yellow, lime and cyan respectively. I27 and linker are in red and green cartoon mode respectively.

## References

1. Nilsson OB, Hedman R, Marino J, Wickles S, Bischoff L, Johansson M, Muller-Lucks A, Trovato F, Puglisi JD, O’Brien EP, Beckmann R, & von Heijne G (2015) Cotranslational Protein Folding inside the Ribosome Exit Tunnel. Cell Rep 12(10): 1533–1540.

2. Nilsson OB, Nickson AA, Hollins JJ, Wickles S, Steward A, Beckmann R, von Heijne G, & Clarke J (2017) Cotranslational folding of spectrin domains via partially structured states. Nat Struct Mol Biol 24(3):221–225.

3. Nilsson OB, Muller-Lucks A, Kramer G, Bukau B, & von Heijne G (2016) Trigger Factor Reduces the Force Exerted on the Nascent Chain by a Cotranslationally Folding Protein. J Mol Biol 428(6):1356–1364.

4. Anfinsen CB (1973) Principles that govern the folding of protein chains. Science 181(4096):223–230.

5. Onuchic JN, Luthey-Schulten Z, & Wolynes PG (1997) Theory of protein folding: the energy landscape perspective. Annu Rev Phys Chem 48(1):545–600.

6. Dill KA & Chan HS (1997) From Levinthal to pathways to funnels. Nat Struct Biol 4(1): 10–19.

7. Tycko R (2004) Progress towards a molecular-level structural understanding of amyloid fibrils. Curr Opin Struct Biol 14(1):96–103.

8. Clark PL (2004) Protein folding in the cell: reshaping the folding funnel. Trends Biochem Sci 29(10):527–534.

9. Dobson CM (2001) Protein folding and its links with human disease. Biochem Soc Symp (68):1–26.

10. Gloge F, Becker AH, Kramer G, & Bukau B (2014) Co-translational mechanisms of protein maturation. Curr Opin Struct Biol 24:24–33.

11. Holtkamp W, Kokic G, Jager M, Mittelstaet J, Komar AA, & Rodnina MV (2015) Cotranslational protein folding on the ribosome monitored in real time. Science 350(6264): 1104–1107.

12. Thommen M, Holtkamp W, & Rodnina MV (2017) Co-translational protein folding:progress and methods. Curr Opin Struct Biol 42:83–89.

13. Krobath H, Shakhnovich EI, & Faísca PF (2013) Structural and energetic determinants of co-translational folding. J Chem Phys 138(21):215–101.

14. Tanaka T, Hori N, & Takada S (2015) How co-translational folding of multi-domain protein is affected by elongation schedule: molecular simulations. PLOS Comput Biol 11(7):e1004356.

15. Wruck F, Katranidis A, Nierhaus KH, Buldt G, & Hegner M (2017) Translation and folding of single proteins in real time. Proc Natl Acad Sci U S A 114(22):E4399–E4407.

16. Jacobs WM & Shakhnovich EI (2017) Evidence of evolutionary selection for cotranslational folding. Proc Natl Acad Sci U S A 114(43): 11434–11439.

17. Samelson AJ, Jensen MK, Soto RA, Cate JH, & Marqusee S (2016) Quantitative determination of ribosome nascent chain stability. Proc Natl Acad Sci U S A 113(47): 13402–13407.

18. O’Brien EP, Christodoulou J, Vendruscolo M, & Dobson CM (2011) New scenarios of protein folding can occur on the ribosome. J Am Chem Soc 133(3):513–526.

19. Knight AM, Culviner PH, Kurt-Yilmaz N, Zou T, Ozkan SB, & Cavagnero S (2013) Electrostatic effect of the ribosomal surface on nascent polypeptide dynamics. ACS Chem Biol 8(6):1195–1204.

20. Cabrita LD, Cassaignau AME, Launay HMM, Waudby CA, Wlodarski T, Camilloni C, Karyadi ME, Robertson AL, Wang X, Wentink AS, Goodsell L, Woolhead CA,Vendruscolo M, Dobson CM, & Christodoulou J (2016) A structural ensemble of aribosome-nascent chain complex during cotranslational protein folding. Nat Struct Mol Biol 23(4):278–285.

21. Fedorov AN & Baldwin TO (1995) Contribution of cotranslational folding to the rate of formation of native protein structure. Proc Natl Acad Sci U S A 92(4): 1227–1231.

22. Nicola AV, Chen W, & Helenius A (1999) Co-translational folding of an alphavirus capsid protein in the cytosol of living cells. Nat Cell Biol 1(6):341–345.

23. Ugrinov KG & Clark PL (2010) Cotranslational folding increases GFP folding yield. Biophys J 98(7):1312–1320.

24. Evans MS, Clarke TF, & Clark PL (2005) Conformations of co-translational folding intermediates. Protein Pept Lett 12(2): 189–195.

25. Goldman DH, Kaiser CM, Milin A, Righini M, Tinoco I, Jr., & Bustamante C (2015) Mechanical force releases nascent chain-mediated ribosome arrest in vitro and in vivo. Science 348(6233):457–460.

26. Fowler SB & Clarke J (2001) Mapping the folding pathway of an immunoglobulin domain: structural detail from Phi value analysis and movement of the transition state. Structure 9(5):355–366.

27. Fowler SB, Best RB, Toca Herrera JL, Rutherford TJ, Steward A, Paci E, Karplus M,& Clarke J (2002) Mechanical unfolding of a titin Ig domain: structure of unfolding intermediate revealed by combining AFM, molecular dynamics simulations, NMRand protein engineering. J Mol Biol 322(4):841–849.

28. Ismail N, Hedman R, Schiller N, & von Heijne G (2012) A biphasic pulling force acts on transmembrane helices during translocon-mediated membrane integration. Nat Struct Mol Biol 19(10):1018–1022.

29. Farias-Rico JA, Selin FR, Myronidi I, & Von Heijne G (2018) Effects of protein size,thermodynamic stability, and net charge on cotranslational folding on the ribosome. bioRxiv:303784.

30. Marino J, Heijne G, & Beckmann R (2016) Small protein domains fold inside the ribosome exit tunnel. FEBS letters 590(5):655–660.

31. Kramer G, Rauch T, Rist W, Vorderwulbecke S, Patzelt H, Schulze-Specking A, Ban N, Deuerling E, & Bukau B (2002) L23 protein functions as a chaperone docking site on the ribosome. Nature 419(6903):171–174.

32. Gong F & Yanofsky C (2003) A transcriptional pause synchronizes translation with transcription in the tryptophanase operon leader region. J Bacteriol 185(21):6472–6476.

33. Seidelt B, Innis CA, Wilson DN, Gartmann M, Armache JP, Villa E, Trabuco LG,Becker T, Mielke T, Schulten K, Steitz TA, & Beckmann R (2009) Structural insight into nascent polypeptide chain-mediated translational stalling. Science 326(5958): 1412–1415.

34. Bischoff L, Berninghausen O, & Beckmann R (2014) Molecular basis for the ribosome functioning as an L-tryptophan sensor. Cell Rep 9(2):469–475.

35. Improta S, Politou AS, & Pastore A (1996) Immunoglobulin-like modules from titinI-band: extensible components of muscle elasticity. Structure 4(3):323–337.

36. Dunkle JA, Xiong L, Mankin AS, & Cate JH (2010) Structures of the Escherichia coli ribosome with antibiotics bound near the peptidyl transferase center explain spectra of drug action. Proc Natl Acad Sci U S A 107(40): 17152–17157.

37. Karanicolas J & Brooks CL, 3rd (2003) Improved Go-like models demonstrate the robustness of protein folding mechanisms towards non-native interactions. J Mol Biol 334(2):309–325.

38. Karanicolas J & Brooks CL, 3rd (2003) The importance of explicit chain representation in protein folding models: an examination of Ising-like models. Proteins 53(3):740–747.

39. Karanicolas J & Brooks CL, 3rd (2003) The structural basis for biphasic kinetics inthe folding of the WW domain from a formin-binding protein: lessons for proteindesign? Proc Natl Acad Sci U S A 100(7):3954–3959.

40. Elcock AH (2006) Molecular simulations of cotranslational protein folding: fragment stabilities, folding cooperativity, and trapping in the ribosome. PLOS Comput Biol 2(7):e98.

41. Muto H, Nakatogawa H, & Ito K (2006) Genetically encoded but nonpolypeptide prolyl-tRNA functions in the A site for SecM-mediated ribosomal stall. Mol Cell 22(4):545–552.

42. Bell GI (1978) Models for the specific adhesion of cells to cells. Science 200(4342):618–627.

43. Ferbitz L, Maier T, Patzelt H, Bukau B, Deuerling E, & Ban N (2004) Trigger factor in complex with the ribosome forms a molecular cradle for nascent proteins. Nature 431(7008):590.

44. Wild K, Halic M, Sinning I, & Beckmann R (2004) SRP meets the ribosome. Nat Struct Mol Biol 11(11): 1049.

45. Frauenfeld J, Gumbart J, Van Der Sluis EO, Funes S, Gartmann M, Beatrix B, Mielke T, Berninghausen O, Becker T, & Schulten K (2011) Cryo-EM structure of the ribosome-SecYE complex in the membrane environment. Nat Struct Mol Biol 18(5):614.

46. Best RB & Hummer G (2016) Microscopic interpretation of folding μ-values usingthe transition path ensemble. Proc Natl Acad Sci U S A 113(12):3263–3268.

47. Sánchez IE & Kiefhaber T (2003) Origin of unusual φ-values in protein folding:evidence against specific nucleation sites. J Mol Biol 334(5):1077–1085.

48. Best RB, Hummer G, & Eaton WA (2013) Native contacts determine protein foldingmechanisms in atomistic simulations. Proc Natl Acad Sci U S A 110(44):17874–17879.

49. Best RB, Fowler SB, Herrera JL, Steward A, Paci E, & Clarke J (2003) Mechanicalunfolding of a titin Ig domain: structure of transition state revealed by combiningatomic force microscopy, protein engineering and molecular dynamics simulations. J Mol Biol 330(4):867–877.

50. Scott KA, Steward A, Fowler SB, & Clarke J (2002) Titin; a multidomain protein thatbehaves as the sum of its parts. J Mol Biol 315(4):819–829.

51. Williams PM, Fowler SB, Best RB, Toca-Herrera JL, Scott KA, Steward A, & Clarke J (2003) Hidden complexity in the mechanical properties of titin. Nature 422(6930):446–449.

52. Wright CF, Lindorff-Larsen K, Randles LG, & Clarke J (2003) Parallel protein-unfolding pathways revealed and mapped. Nat Struct Biol 10(8):658–662.

53. Wright CF, Steward A, & Clarke J (2004) Thermodynamic characterisation of twotransition states along parallel protein folding pathways. J Mol Biol 338(3):445–451.

54. Borgia MB, Nickson AA, Clarke J, & Hounslow MJ (2013) A mechanistic model foramorphous protein aggregation of immunoglobulin-like domains. J Am Chem Soc 135(17):6456–6464.

55. Botello E, Harris NC, Sargent J, Chen WH, Lin KJ, & Kiang CH (2009) Temperatureand chemical denaturant dependence of forced unfolding of titin I27. JPhys Chem B 113(31): 10845–10848.

56. Chen H, Yuan G, Winardhi RS, Yao M, Popa I, Fernandez JM, & Yan J (2015) Dynamics of equilibrium folding and unfolding transitions of titin immunoglobulindomain under constant forces. J Am Chem Soc 137(10):3540–3546.

57. Lu H, Isralewitz B, Krammer A, Vogel V, & Schulten K (1998) Unfolding of titinimmunoglobulin domains by steered molecular dynamics simulation. Biophys J 75(2):662–671.

58. Nunes JM, Mayer-Hartl M, Hartl FU, & Muller DJ (2015) Action of the Hsp70chaperone system observed with single proteins. Nat Commun 6:6307.

59. Yagawa K, Yamano K, Oguro T, Maeda M, Sato T, Momose T, Kawano S, & Endo T(2010) Structural basis for unfolding pathway-dependent stability of proteins:vectorial unfolding versus global unfolding. Protein Sci 19(4):693–702.

60. Zheng W, Schafer NP, & Wolynes PG (2013) Frustration in the energy landscapes ofmultidomain protein misfolding. Proceedings of the National Academy of Sciences 110(5): 1680–1685.

61. Carrion-Vazquez M, Oberhauser AF, Fowler SB, Marszalek PE, Broedel SE, Clarke J, & Fernandez JM (1999) Mechanical and chemical unfolding of a single protein: acomparison. Proc Natl Acad Sci U S A 96(7):3694–3699.

62. Su T, Cheng J, Sohmen D, Hedman R, Berninghausen O, von Heijne G, Wilson DN,& Beckmann R (2017) The force-sensing peptide VemP employs extreme compactionand secondary structure formation to induce ribosomal stalling. eLife 6:e25642.

63. Ziv G, Haran G, & Thirumalai D (2005) Ribosome exit tunnel can entropicallystabilize a-helices. Proc Natl Acad Sci US A 102(52):18956–18961.

64. Guinn EJ, Tian P, Shin M, Best RB, & Marqusee S (2018) A small single-domain protein folds through the same pathway on-and off-the ribosome. bioRxiv:347864.

65. Mercier E & Rodnina MV (2018) Co-translational Folding Trajectory of the HemK Helical Domain. Biochemistry.

66. Samelson AJ, Bolin E, Costello SM, Sharma AK, O’Brien EP, & Marqusee S (2018) Kinetic and structural comparison of a protein’s cotranslational folding and refolding pathways. Sci Adv 4(5):eaas9098.

67. Evans MS, Sander IM, & Clark PL (2008) Cotranslational folding promotes β-helix formation and avoids aggregation in vivo. J Mol Biol 383(3):683–692.

68. Mirzadeh K, Martinez V, Toddo S, Guntur S, Herrgard MJ, Elofsson A, Norholm MH, & Daley DO (2015) Enhanced Protein Production in Escherichia coli by Optimization of Cloning Scars at the Vector-Coding Sequence Junction. ACS Synth Biol 4(9):959–965.

69. Li X, Mooney P, Zheng S, Booth CR, Braunfeld MB, Gubbens S, Agard DA, & Cheng Y (2013) Electron counting and beam-induced motion correction enable near-atomic-resolution single-particle cryo-EM. Nat Methods 10(6):584–590.

70. Scheres SH (2012) RELION: implementation of a Bayesian approach to cryo-EM structure determination. J Struct Biol 180(3):519–530.

71. Zhang K (2016) Gctf: Real-time CTF determination and correction. J Struct Biol 193(1): 1–12.

72. Kucukelbir A, Sigworth FJ, & Tagare HD (2014) Quantifying the local resolution of cryo-EM density maps. Nat Methods 11(1):63–65.

73. Pettersen EF, Goddard TD, Huang CC, Couch GS, Greenblatt DM, Meng EC, & Ferrin TE (2004) UCSF Chimera--alysis. J Comput Chem 25(13): 1605–1612.

74. Hess B, Kutzner C, Van Der Spoel D, & Lindahl E (2008) GROMACS 4: algorithms for highly efficient, load-balanced, and scalable molecular simulation. J Chem Theory and Comput 4(3):435–447.

75. Hess B, Bekker H, Berendsen HJ, & Fraaije JG (1997) LINCS: a linear constraint solver for molecular simulations. J Comput Chem 18(12): 1463–1472.

76. Sirur A, Knott M, & Best RB (2014) Effect of interactions with the chaperonin cavity on protein folding and misfolding. Phys Chem Chem Phys 16(14):6358–6366.

77. Bortz AB, Kalos MH, & Lebowitz JL (1975) A new algorithm for Monte Carlo simulation of Ising spin systems. J Comput Phys 17(1): 10–18.

78. Öhman A, Öman T, & Oliveberg M (2010) Solution structures and backbone dynamics of the ribosomal protein S6 and its permutant P54-55. Protein Sci 19(1): 183–189.

79. Pascual J, Pfuhl M, Walther D, Saraste M, & Nilges M (1997) Solution structure of the spectrin repeat: a left-handed antiparallel triple-helical coiled-coil1. J Mol Biol 273(3):740–751.

